# msqrob2TMT: robust linear mixed models for inferring differential abundant proteins in labelled experiments with arbitrarily complex design

**DOI:** 10.1101/2024.03.29.587218

**Authors:** Stijn Vandenbulcke, Christophe Vanderaa, Oliver Crook, Lennart Martens, Lieven Clement

**Affiliations:** VIB-UGent Center for Medical Biotechnology, VIB, Ghent, Belgium; Department of Biomolecular Medicine, Ghent University, Ghent, Belgium; Department of Applied Mathematics, Computer Science and Statistics, Ghent University, Ghent, Belgium; Department of Statistics, University of Oxford, Oxford, UK

## Abstract

Labelling strategies in mass spectrometry (MS)-based proteomics enable increased sample throughput by acquiring multiplexed samples in a single run. However, contemporary designs often require the acquisition of multiple runs, leading to a complex correlation structure. Addressing this correlation is key for correct statistical inference and reliable biomarker discovery. Therefore, we present msqrob2TMT, a set of mixed model-based workflows tailored toward differential abundance analysis for labelled MS-based proteomics data. Thanks to its increased flexibility, msqrob2TMT can model both sample-specific and feature-specific (e.g. peptide or protein) covariates, which unlocks the inference to experiments with arbitrarily complex designs as well as to correct explicitly for feature-specific properties. We benchmark our novel workflows against the state-of-the-art tools MSstatsTMT and DeqMS in a spike-in study. We show that our workflows are modular, more flexible and have improved performance by adopting robust ridge regression. We also found that reference channel normalization and imputation can have a deleterious impact on the statistical outcome. Finally, we demonstrate the significance of msqrob2TMT on a real-life mice study, showcasing the importance of effectively accounting for the hierarchical correlation structure in the data.

## 1 Introduction

High-throughput LC-MS-based proteomic workflows are widely used to quantify differential protein abundance between samples. Relative protein quantification is typically done using label-free or labelled workflows. The latter can be achieved by stable isotope labelling workflows such as metabolic and post metabolic labelling, which gained a lot of traction over the last years. As opposed to label-free workflows, labelling has the advantage that it avoids run-to-run differences in the identification and quantification of peptides by pooling and analyzing multiple samples in a single run. This allows researchers to compare the proteomes of multiple conditions or treatments in a single MS-run, providing a more comprehensive view of the proteome and increasing the statistical power of the analysis. Indeed, with the current tandem mass tags (TMT) kits up to 18 samples can be multiplexed in the same run (Li et al. 2021). However, contemporary experiments typically involve more complex designs for which the samples often no-longer fit on a single run.

This has far reaching consequences for the downstream data analysis. Indeed, labelled experiments with multiple MS runs typically involve distinct biological replicates on distinct runs and often multiple technical MS runs. Similar to unlabelled approaches, they again suffer from many missing peptide intensities between MS runs (Brenes et al. 2019). Moreover, these designs also implies a complex correlation structure with within and between run variance components, and possibly correlation between technical repeats from the same biological repeat. Moreover, in large experiments these complex designs are often unbalanced and require flexible models that can address this hierarchical correlation structure. There are few bespoke software tools that can handle these complex correlation structure. MSstatsTMT (Huang et al. 2020b) is one of the notable exceptions, which has been specifically designed for experiments with multiple conditions, multiple biological replicate runs, multiple technical replicate runs, and unbalanced designs.

In this publication we develop a custom data analysis workflow for labelled MS-based proteomics experiments in our msqrob2 universe in R/Bioconductor. msqrob2 has been originally developed for label free workflows, where it has been shown to have advantages over other state-of-the-art tools both in terms of its performance as well as by its modular and more transparent implementation (Sticker et al. 2020). Here we will bring these advantages to the analysis of labelled experiments. Similar to MSstatsTMT, our workflows will address the hierarchical correlation structure in contemporary labelled MS-based proteomics experiments by building upon mixed models. In contrast to MSstatsTMT, however, we will avoid imputation of missing PSM intensities and assume missingness at random after accounting for peptide effects. Nonetheless, users can still rely on the default imputation strategies of QFeatures or implement custom functions needed for their specific application. Moreover, msqrob2 also provides ridge regression and robust M-estimation, which can further stabilise the parameter estimation.

We show that our novel msqrob2TMT workflows outperform state-of-the-art tools by bench-marking them to DEqMS (Zhu et al. 2020) and MsStatsTMT (Huang et al. 2020b) using spike-in and real data. Note that we did not include Proteome Discoverer in our benchmark as we use the same datasets as in the original MSstatsTMT publication, which already showed superior performance of MSstatsTMT (Huang et al. 2020b). Importantly, our msqrob2TMT workflows for labelled experiments also provide researchers with full flexibility to develop both peptide-based as well as protein-level data analysis workflows with custom pre-processing steps and user-specified models tailored toward their specific applications. The newest release unlocked more complex designs by providing full compatibility with the model specification of the popular lme4 R package for mixed models, and adding the flexibility to use both feature level and sample level covariates. The former allows msqrob2 and msqrob2TMT to model longitudinal and clustered designs. The latter is provided by building on the QFeatures (Gatto and Vanderaa 2022) data infrastructure, with which we also guarantee that the input data are never lost, but remains available and is linked to the pre-processed, normalised and summarised assays as well as to the model output, insuring transparency, trace-ability and reproducibility.

## 2 Materials and Methods

In this section we first introduce the datasets that are used to evaluate our workflows and to benchmark them to state-of-the-art methods. Next, we introduce the preprocessing. We continue with focusing on modelling different sources of variation in labelled proteomics experiments. Then, we introduce our novel msqrob2TMT workflows and these of MSstatsTMT and DEqMS. We conclude this section with the different metrics to benchmark the performance.

### 2.1 Datasets

Two datasets are used to evaluate and benchmark the different tools. A spike-in dataset and real mouse case study.

#### 2.1.1 Spike-in Dataset

The spike-in dataset was taken from ProteomeXchange (PXD0015258). It consists of controlled mixtures with known ground truth. UPS1 peptides at concentrations of 500, 333, 250, and 62.5 fmol were spiked into 50 g of SILAC HeLa peptides, each in duplicate. These concentrations form a dilution series of 1, 0.667, 0.5, and 0.125 relative to the highest UPS1 peptide amount (500 fmol). A reference sample was created by combining the diluted UPS1 peptide samples with 50g of SILAC HeLa peptides. All dilutions and the reference sample were processed in duplicate, resulting in a total of ten samples. These samples were then treated with TMT10-plex reagents and combined before LC-MS/MS analysis. This protocol was repeated five times, each with three technical replicates, totaling 15 MS runs.

#### 2.1.2 Mouse dataset

The mouse dataset was taen from ProteomeXchange (PXD005953). (Plubell et al. 2017) split twenty mice into groups for a study on the impact of low or high fat diets. The mice were split into groups based on their diet, either low-fat (LF) or high-fat (HF), and the duration of the diet, which was classified as short (8 weeks) or long (18 weeks). Four separate groups were formed, each consisting of five mice, based on the combination of diet type and duration. Samples from the epididymal adipose tissue of these mice were then randomly distributed across three TMT 10-plex mixtures for analysis. In each mixture, two reference channels were used, each containing pooled samples that included a range of peptides from all the samples. Not all channels were used, leading to an unbalanced design. Each TMT mixture was divided into eight parts and subjected to SPS, resulting in a total of 27 MS runs.

### 2.2 Pre-processing

The data analysis starts by log2 transforming the measured intensities. Next, we introduce an additional filtering step because two PSMs can have identical intensity values due to the Mascot search nodes. Indeed, one search is done on the SwissProt protein database and the other search is done on a Sigma UPS protein database. This was done to find which PSMs belonged to the HeLa cells, which were metabolically labelled, as well as metabolically labelled spiked-in UPS proteins. The identification, however, is not perfect as some PSMs are matched in both nodes, which leads to duplicated PSMs with identical intensity values. We remove these PSMs as it is not possible to determine if they match the spiked-in or the background UPS proteins. We further remove PSMs that have less than 6 intensity values, this cutoff is arbitrarily chosen. In the case of multiple spectra for the same PSMs, the PSM with the highest sum of intensity values is chosen and was used here for comparison reasons. The pre-processing then diverges for the three methods tested in this work.

#### 2.2.1 msqrob2TMT

The msqrob2TMT workflow proceeds with the normalization of the TMT channels by subtracting the median log2 intensity of each channel to account for loading differences. Hence, the normalization assumes that the large majority of the proteins are not differentially abundant. For our protein-level workflows, the PSMs are summarised into protein abundance values by adopting Tukey’s median polish method to the PSM data of each protein in each run, separately. It robustly estimates the parameters of the following additive model for each protein within a run. Note, that we drop the protein index for notational convenience:

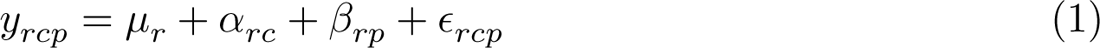

with *y*_*rcp*_ the log-transformed and normalised intensity for PSM *p* in run *r* and channel *c*, μ_*r*_ is the overall run effect, α_*rc*_ is the channel effect nested within run and β_*rp*_ the spectrum effect and ∈_*rcp*_ the residual error. The channel effects for each protein nested in a run α_*rc*_ are used as the summarised abundance values. No imputation is done in our default msqrob2TMT workflows, although users can rely on their custom imputation strategies or use the default functionalities in the QFeatures package.

#### 2.2.2 MSstatsTMT

The default MSstatsTMT workflow proceeds with equalizing the medians across PSMs, channels and runs. The missing values are imputed using an accelerated failure time model. After imputation, PSM intensities are summarised to protein intensities by Tukey’s median polish. Finally, protein intensities are reference normalised by subtracting the median of the corresponding protein intensities of the reference channels.

#### 2.2.3 DEqMS

The DEqMS workflow proceeds with the summarization of the PSM intensities into protein intensities by adopting the median sweep algorithm to the PSM data of each protein in each run, separately. Note, that it can be considered as a one-step median polish. Indeed, log_2_ PSM intensities are first centered by substracting the corresponding median of log2 PSM intensities in their corresponding spectrum. Next, the spectrum centered PSM intensities in each TMT channel for each protein are summarised into protein expression values by median aggregation. Finally, the protein summaries are normalised by subtracting the median of all summarised protein expression values in their corresponding channel.

### 2.3 Modelling different sources of variation

Proteomics data contain several sources of variation that need to be accounted for by the model. First, there is the variation of interest induced by the experimental treatment. We model this source of variation as a fixed effect, which we consider non-random, i.e. the treatment effect is assumed to be the same in repeated experiments, but it is unknown and has to be estimated. When modelling a typical label-free experiment at the protein level, the model boils down to a linear model, again we suppress the index for protein:

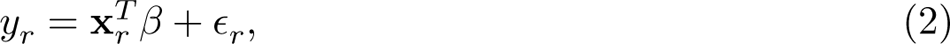

with *y*_*r*_ the log2-normalised protein intensity in run r; **x**_*r*_ a vector with the covariate pattern for the sample in run r encoding the treatment, potential batch effects and confounders; β_*r*_ the vector of parameters that model the association between the covariates and the outcome; and ∈_*r*_ the residuals reflecting variation that is not captured by the fixed effects. Note that **x**_*r*_ allows for a flexible parameterization of the treatment beyond a single covariate, including continuous and categorical variables as well as their interactions. For all models considered in this work, we assume the residuals to be independent and identically distributed (i.i.d) according to a normal distribution with zero mean and constant variance, i.e. ∈_*r*_ ∼ *N*(0, σ^2^), that can differ from protein to protein.

Since label-free experiments contain only a single sample per run, run-specific effects will be absorbed in the residuals. However, the data analysis of labelled experiments, e.g. using TMT multiplexing, involving multiple MS runs has to account for run- and label-specific effects, explicitly. Hence, model (2) is extended to:

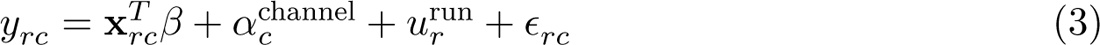

with *y*_*rc*_ the normalised log_2_ protein intensities in run *r* and channel *c*, α^channel^ the effect introduced by the label of channel *c*, and *u*^run^ the effect for MS run *r*.

If all treatments are present in each run, and if channel swaps are performed so as to avoid confounding between channel and treatment, then the model parameters can be estimated using fixed channel and run effects. Indeed, for these designs run acts as a blocking variable as all treatment effects can be estimated within run. However, for more complex designs this is no longer possible and the estimation of the mean model parameters can involve both within and between run variability. For these designs we can resort to mixed models where the run effect is modelled using a random effect, i.e. it is considered as a random sample from the population of all possible runs, which are assumed to be i.i.d normally distributed with mean 0 and constant variance, *u*_*r*_ ∼ *N*(0, σ^2,run^). The use of random effects thus models the correlation in the data, explicitly. Indeed, protein intensities that are measured within the same run will be more similar than protein intensities between runs.

Some experiments also include technical replication where a TMT mixture can be acquired multiple times. This again will induce correlation. Indeed, protein intensities from the same mixture will be more alike than those of different mixtures. Hence, we also include a random effect to account for this pseudoreplication, i.e. *u*^mix^ ∼ *N*(0, σ^2,mix^). The model thus extends to:

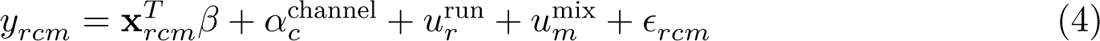

with *m* the index for mixture.

Above, we modelled the data at the protein level. However, we could also directly estimate the treatment effect from PSM-level data. This, again will induce additional levels of correlation. Indeed, the intensities for the different reporter ions in a TMT run within the same spectrum (PSM) will be more similar than intensities between PSMs. We therefore need to add an additional random effect term to account for the within PSM correlation structure, i.e. *u*^PSM^ ∼ *N*(0, σ^2,PSM^). Moreover, each sample now contains multiple PSM intensities for each protein. Hence, intensities from different PSMs for a protein in the same sample will be more alike than intensities of different PSMs for the same protein between samples, and we will address this correlation with a sample specific random effect. Note, that each sample is uniquely defined by its run and channel index. Hence, we can parameterise the sample effect using a random channel effect nested in run, i.e. *u*^*c*ℎchannel^ ∼ *N*(0, σ^2,channel^). The model then becomes:

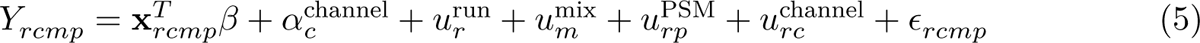

with *y*_*rcmp*_ the log2-normalised PSM intensities for run *r* with label *c* in mixture *m* and PSM *p*. Note, that the PSM random effect is also nested within each run since each spectrum is described by run-specific characteristics.

The fixed channel effect, α^channel^, that models the effect of the isobaric label is often ignored and we will drop it from the model in the remainder of the paper.

### 2.4 msqrob2TMT workflows

Our PSM-level msqrob2TMT workflows start from PSM-level data, and our protein-level msqrob2TMT workflows start from summarised data upon median polish summarization model (1). msqrob2TMT fits linear mixed models for each protein which can be implemented in msqrob2 version 1.10 from Bioconductor version 3.18 (Huber et al. 2015). If there is not enough data to estimate all model parameters, then msqrob2 returns a “FitError”. Indeed, missing data can change the parameterisation of the fixed effects and we feel that an intervention of a skilled data analyst is needed to ensure that the correct research hypothesis is assessed for these proteins.

The treatment effects can be regularised using ridge penalization by exploiting the link between ridge regression and mixed models, and outliers can be accounted for using M-estimation with Huber weights (Goeminne et al. 2020). Upon estimation, fold changes can be calculated using linear combinations of the mean model, i.e. contrasts on the parameters for the covariates related to the treatment. The variance components are estimated using REstricted Maximum Likelihood (REML) and the variance of the errors is stabilised by borrowing strength across proteins using the default empirical Bayes step of msqrob2. Statistical inference on the contrasts of interest is done using Wald tests.

Note, that our msqrob2 framework is very flexible. From version 1.10 onwards the model specification is fully compatible with the lme4 R package for fitting mixed models. Moreover, it also exploit the QFeatures data infrastructure to enable the use of both feature-level (rowData) variables as well as sample-level (colData) variables in the model, unlocking the analysis to infer and correct for both feature and sample specific covariates, and their interactions. Hence, our current msqrob2 implementation can therefore be used for arbitrarily complex designs as long as they can be specified within the linear mixed model framework.

### 2.5 MSstatsTMT

MSstatsTMT fits linear mixed models for each protein and is performed using MSstatsTMT version 2.10 (Huang et al. 2020a) from Bioconductor version 1.18. The variance components are also estimated using REML. Note, however, that MSstatsTMT can only be applied to experiments with a treatment that can be modelled using one factor and random effects for run and mixture as opposed to msqrob2, which allows the user to specify models for arbitrarly complex designs. If there is not enough data to estimate the parameters of the full model, MSstatsTMT will reduce the model to perform inference on as many proteins as possible.

### 2.6 DEqMS

DEqMS uses conventional linear model for each protein and is performed using the DEqMS version 1.20 (Zhu 2023) from Bioconductor version 1.18. DEqMS cannot model random effects. Therefore, it can only address run using a fixed block effect and only provides valid inference for treatment effects that can be estimated within run. Hence, it also cannot address technical replication.

### 2.7 Models for spike-in and case study

We first introduce the models for the spike-in study and then for the mouse case study.

#### 2.7.1 Spike-in study

For the spike-in data, the treatment of interest is the UPS1 dilution, which can be modelled as a continuous variable. However, MSstatsTMT is mostly used to assess anova designs with one factor, which can be adopted here by considering the UPS1 dilution in the spike-in study as a categorical variable with 4 levels (A: 0.125, B: 0.5, C: 0.667 and D:1). Hence, **x**_*r*_ is parameterised using the lowest spike-in condition A as a reference group and dummy variables *x*^*B*^, *x*^*C*^ and *x*^*D*^ for each of the remaining spike-in conditions (B-D), with *x*_*j*_ = 1 if the sample belongs to the spike-in condition *j* and zero otherwise.

Another constraint is that DEqMS cannot estimate random effects and hence it cannot address technical replication. Therefore, one technical repeat is used to enable the comparison with the DEqMS workflow. For a protein-level analysis with msqrob2, MSstatsTMT and DEqMS, we can reduce the model (4) to

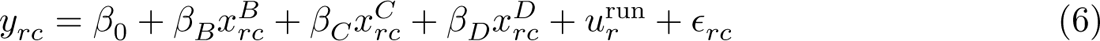

So the parameter vector β consists of an intercept β_0_ and the log_2_ fold changes (FC) log_2_ FC_*B*−*A*_ = β_*B*_ for comparison B vs A, log_2_ FC_*C*−*A*_ = β_*C*_ for C vs A, and log_2_ FC_*D*−*A*_ = β_*D*_ for D vs A. Note, that the remaining fold changes are linear combinations of the model parameters, also referred to as contrasts, e.g. for comparison C vs B it becomes log_2_ FC_*C*−*B*_ = β_*C*_ − β_*B*_. msqrob2TMT is the only workflow that can estimate the PSM-level model (5). Note that we dropped the random mixture effect for modelling the subset of the spike-in study with one technical repeat. Hence model (5) reduces to:

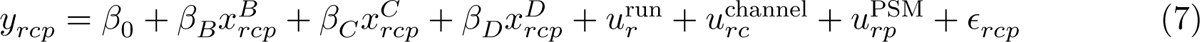

Finally, benchmarking results for msqrob2TMT and MSstatsTMT for the full spike-in study imply model (4) for the protein-level workflow and model (5) for the psm-level workflow.

Note, that we also dropped the fixed effect for channel label α_*c*_ from all models.

#### 2.7.2 Mouse case study

Here, we limit the comparison to the best-performing workflows, msqrob2TMT and MSstatsTMT, as benchmarked on the spike-in study. For the mouse case study, the factor treatment is encoded as diet × duration because MSstatsTMT can only model the treatment effect using one factor. msqrob2TMT also has the flexibility to use the more natural parameterisation with a main effect for diet, a main effect for duration and a diet × duration interaction, which would lead to the same model fit. Again, we drop the fixed effect for channel label α_*c*_.

For the protein-level workflows the following model is used:

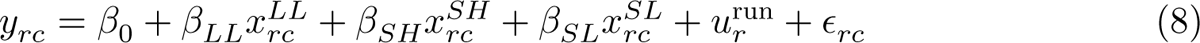

with *x*^*LL*^_rc_, *x*^SH^_rc_ and *x*^SL^_rc_ dummy variables that are 1 if sample *rc* belongs to group LongLF, ShortHF, ShortLF, respectively and zero otherwise. Hence, group LongHF is the reference group.

The msqrob2TMT workflow at PSM level reduces to:

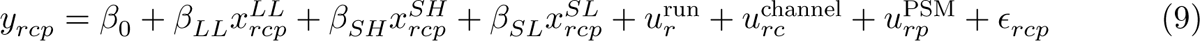

### 2.8 Method performance

The methods are benchmarked using the spike-in study. In particular we use the number of true positive (TP, spiked-in UPS) and false positive proteins (FP, HeLa proteins) that are reported. We also construct true positive rate (TPR) - false discovery proportion (FDP) plots. Note, that TPR is the fraction of the truly DA proteins (spiked-in UPS) picked up by the method and FDP is the fraction of false positives (HeLa) on the total number of proteins flagged as DA. On the TPR-FDP plot we also indicate the observed FDP at 5% FDR cut-off, which is expected to be close to 5%. We also evaluate the log_2_ FC estimates and compare them with the ground truth, which is zero for the non-spiked HeLa background proteins and the log_2_ - ratio of the spike-in dilutions for spiked-in UPS proteins.

To assess the type I error (false positives) in the real mouse study, we also performed a mock analysis and randomly assigned two mock levels within the conditions. Hence, all proteins with a significant mock effect are false positives. The model at protein-level is specified as:

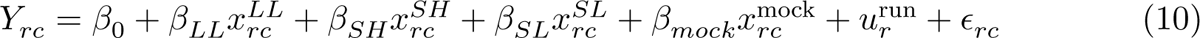

with *x*^mock^ an additional dummy variable that 1 if sample *rc* belongs to mock group 2 and zero otherwise. This is the model for the mice study where we introduced an additional fixed effect for the mock factor. Since this design cannot be recasted in a design with one factor for treatment, this model cannot be estimated with MSstatTMT.

The PSM model for the mock analysis becomes:

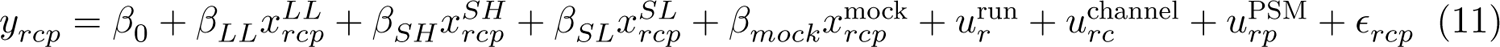

## 3 Results

We will compare our novel msqrob2TMT workflows for labelled proteomics experiments to state-of-the-art methods DEqMS and MSstatsTMT. The different functionalities of the tools can be found in Figure 1. Note, that msqrob2TMT is the only workflow that can be used to model arbitrarily complex designs. Indeed, MSstatsTMT can only model treatment effects that can be specified using a single factor and can only address correlation induced by run and mixture effects, while DEqMS cannot model the correlation in the data with random effects. In this section we first compare the existing TMT workflows of DEqMS and MSstatsTMT to our novel msqrob2TMT workflows and critically assess differences in the performance in a spike-in study. Next, we explain how and why our msqrob2TMT workflows improve upon the state-of-the-art methods. Then, we illustrate that our msqrob2TMT workflows recover more biological relevant proteins in a biological mouse study (PXD00593). We conclude this section with mock analysis where we show that our novel msqrob2TMT workflow also controls the type I error on real data.

**Figure 1:**
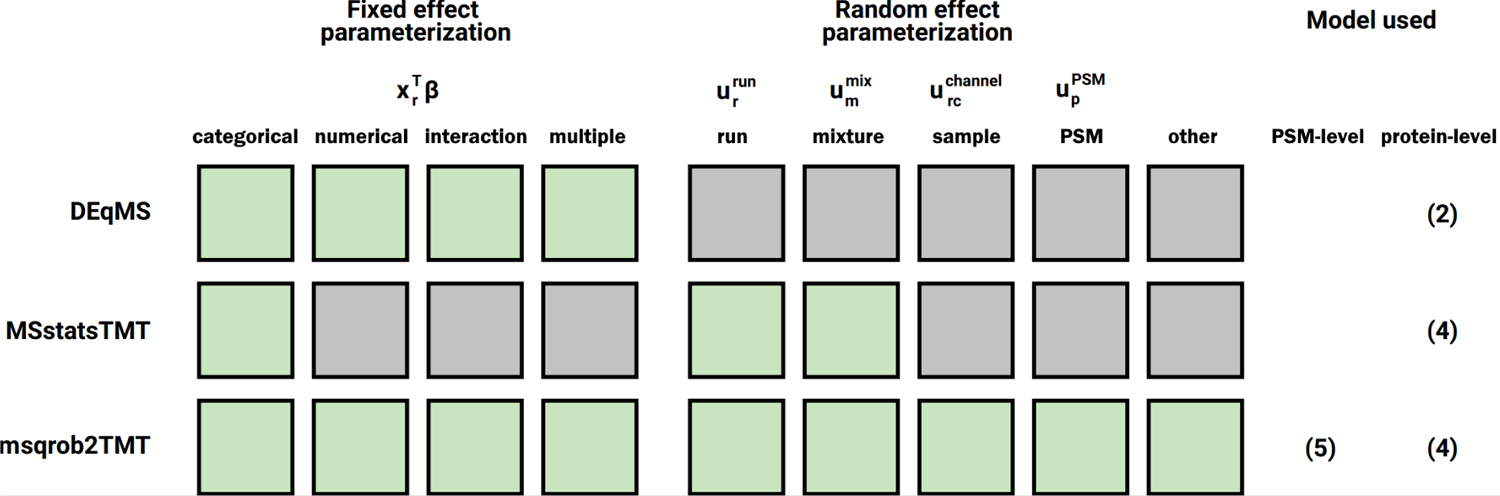
Overview of the tools considered in this work and the model parameterization they enable. Green boxed indicate that the method supports the modelling of the parameter. Note that “other” refers to other random effects implied by the design, e.g. in longitudinal studies. More information on the description of the models and their parameters can be found in the Material and Methods section. Note, that the fixed channel effect in models (4) and (5) were omitted.

### 3.1 Spike-in Study

In this study (PXD0015258) 40 UPS proteins (UPS1 mix) were spiked into the HeLa protein background at four different concentrations. Each spike-in concentration was multiplexed twice in a 10-plex TMT mixtures together with two reference channels. Five 10-plex TMT mixtures were made and each mixture was run in technical triplicate on the mass-spectrometer. We therefore know the ground truth: only the spiked-in UPS proteins are differentially abundant (DA).

We used this spike-in study to compare the performance of the default TMT workflows of DEqMS, MSstats and three msqrob2TMT workflows:

1. msqrob2-rlm (Robust Linear Model) that uses msqrob2’s default workflow that models the summarised protein level expression values using robust linear models that corrects for outliers using M-estimation.
2. msqrob2-rrilmm (Robust RIdge Linear Mixed Model) a novel workflow that models the summarised protein level expression values using a robust linear mixed model that addresses the correlation within TMT-plex using a random effect, regularises the fold change estimates between spike-in conditions using a ridge penalty and corrects for outliers using M-estimation.
3. msqrob2-psm-rrilmm (peptide spectrum match Robust RIdge Linear Mixed Model) a novel workflow that directly models normalised PSM intensities using a robust linear mixed model with random effects for sample, PSM and run, a ridge penalty to regularise the fold change estimates and corrects for outliers using M-estimation.

Note, that the DEqMS and the default msqrob2-rlm workflow cannot address the hierarchical correlation structure in labelled experiments with multiple runs. When all conditions are present in each run, however, they can correctly infer fold changes by introducing a block factor for run. But, they cannot correctly account for technical repeats. We, therefore, assess the performance of the different methods using only one technical repeat for each mixture in the main manuscript and provide the results for the full study in Supplementary Information.

#### 3.1.1 Comparison of the workflows

In Figure 2 we compare the performance of the different methods using True positive rate (TPR)-False discovery proportion (FDP) curves. Note, that the former is the fraction of spiked UPS proteins that are returned as DA and the latter is the ratio of HeLa proteins on the total number of DA proteins that are returned. In Table 1 the number of true positives (TP), i.e. the spiked-in UPS proteins and the false positives (FP), i.e. the HeLa proteins that are returned as DA at the 5% FDR significance level are given.

**Figure 2:**
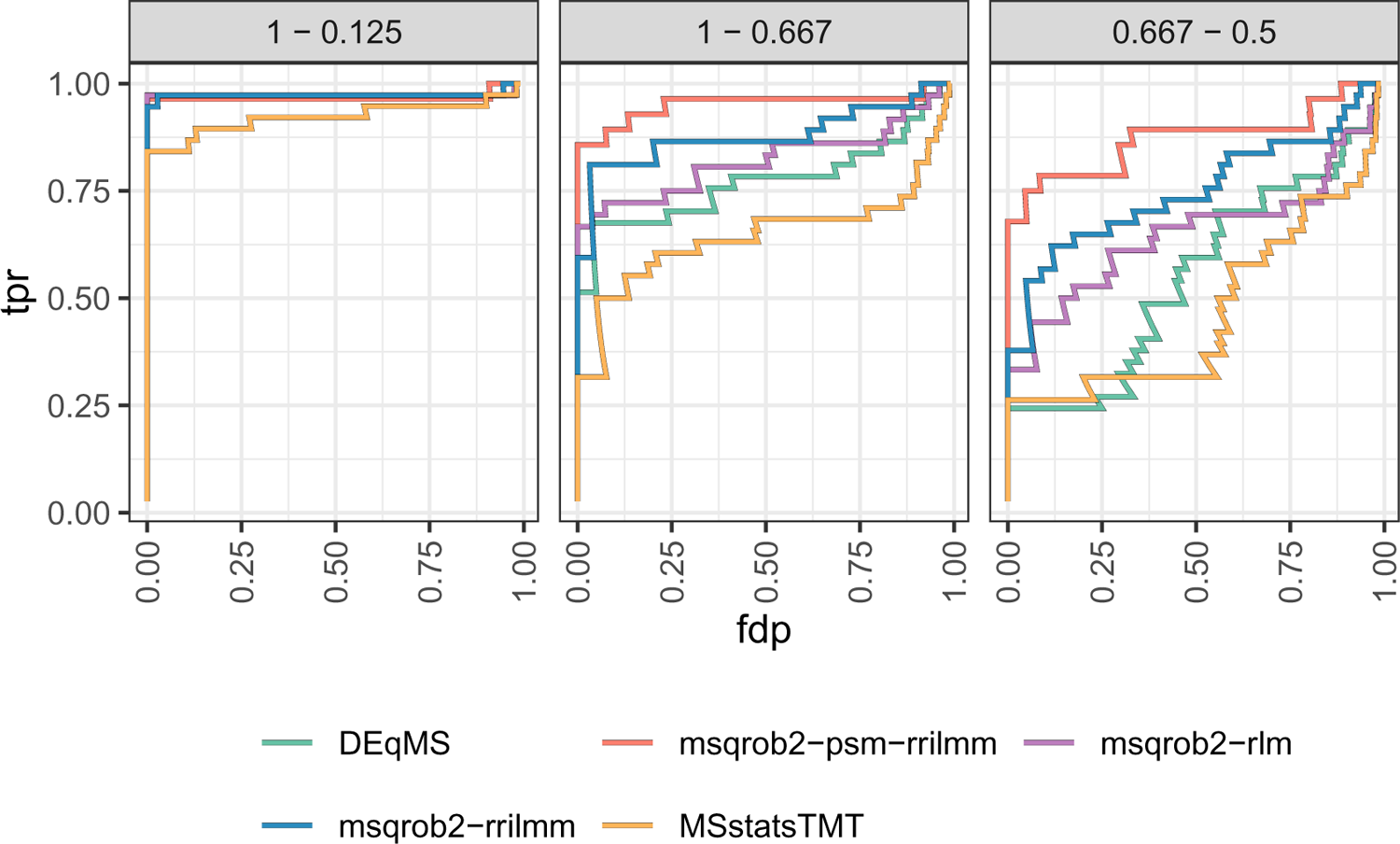
True positive rate (tpr) - false discovery proportion (fdp) plots for DEqMS, msqrob2TMT and MSstatsTMT workflows. Three comparisons of the spike-in dilutions illustrating the differences are shown, see Supplementary Figure: 1 for all the spike-in concentration comparisons.

**Table 1:** True positives (TP) and false positives (FP) at 0.05 FDR adjusted p-value cutoff for the DEqMS, msqrob2TMT and MSstatsTMT workflows. Three spike-in concentration comparisons illustrating the differences are shown, see Supplementary Table: 1 for the comparisons of all spike-in concentrations.

|  | TP | FP | TP | FP | TP | FP |
| --- | --- | --- | --- | --- | --- | --- |
|  | 1 - 0.667 | 1 - 0.667 | 0.667 - 0.5 | 0.667 - 0.5 | 1 - 0.125 | 1 - 0.125 |
| DEqMS | 21 | 1 | 9 | 0 | 36 | 5 |
| MSstatsTMT | 16 | 1 | 5 | 0 | 34 | 5 |
| msqrob2-psm-rilmm | 23 | 0 | 17 | 0 | 27 | 1 |
| msqrob2-rlm | 21 | 0 | 12 | 1 | 35 | 5 |
| msqrob2-rilmm | 22 | 1 | 14 | 0 | 35 | 1 |

Figure 2 and Table 1 show for the most extreme comparison, log2 fold change (FC) of 3 (log 2(1/0.125)), that all methods have a good performance. However, we observed that our msqrob based workflows slightly outperform MSstatsTMT. Moreover, at the 5% FDR DEqMS, msqrob2-rlm and MSstat return more false positives than our novel msqrob2TMT workflows. Indeed, their observed false discovery proportion (FDP = FP/(FP+TP)) is 0.122, 0.125 and 0.128, respectively indicating that they do not control the FDR at the nominal 5% level. For our novel workflows, the FDP is close to the 5%-level, i.e. 0.028 for msqrob2-rrilmm and 0.036 for msqrob2-psm.

For the comparisons involving log2 fold changes larger or equal than two, similar results are observed (see Supplementary Figure: 1).

Our robust ridge regression workflows, however, clearly outperform DEqMS and MSstats for intermediate (log 2(1/0.667) = 0.58), and, low fold changes (log 2(0.667/0.5) = 0.42), which are more difficult to pick up (Figure 2). Indeed, msqrob2-rrilmm and msqrob2-psm return up to three times more spike-in proteins than MSstats and DEqMS at the 5% FDR level, while returning at most 1 false positive. Note, that msqrob2-psm-rrilmm is slightly more sensitive than our dedicated msqrob2-rrilmm workflow that summarises the PSM intensities first at the protein-level. msqrob2-psm-rrilmm, however, has the drawback that it is computationally more complex and that no corresponding protein expression values are returned by the model. It is left to the user to decide how to aggregate them for visualisation.

The full dataset with all technical replicates can only be analysed correctly with our bespoke labelled msqrob workflows and MSstatsTMT. Note, that the results are inline with the analysis in Figure 2, however, the difference in performance is smaller (See Supplementary Figure: 4).

Figure 3 shows that the log2 fold change estimates for non-spiked proteins are unbiased for all methods. However, the log2-FC for our msqrob workflows with ridge regularisation are less variable.

**Figure 3:**
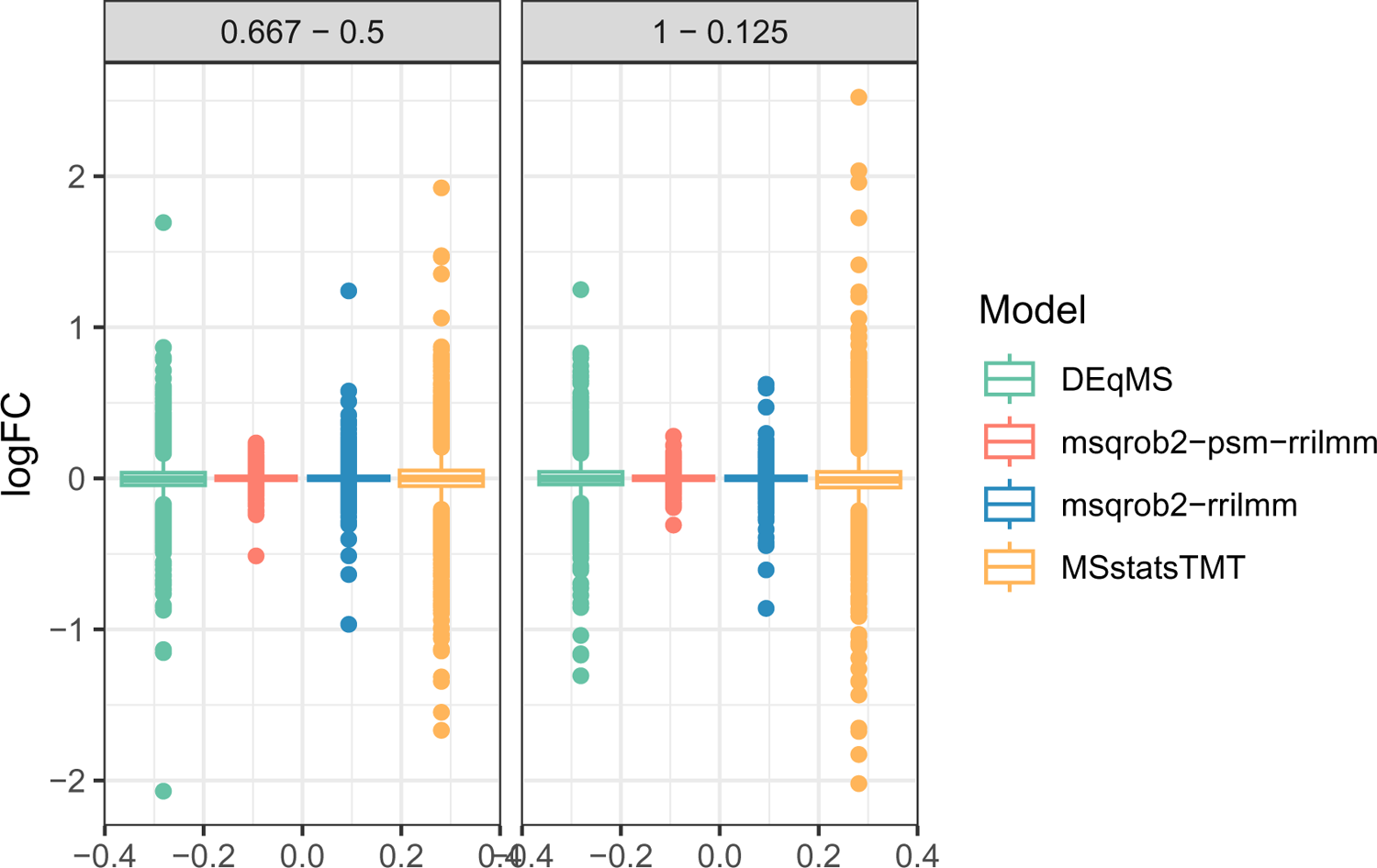
Boxplots showing the logFC distribution of two spike-in concentration comparisons for the non spike-in proteins for the comparisons with the highest and lowest fold change between spike in concentrations estimated by DEqMS, MSstatsTMT and msqrob2TMT workflows. The models using ridge regression show a more narrow boxplot due to the regularization.

Figure 4, shows that the estimated fold changes for spike-in proteins are always underestimated for all methods. This has been reported in the literature and is probably due to interference of cofragmentation of peptide ions which leads to underestimation of the fold changes between the reporter ions (Savitski et al. 2011; Ow et al. 2009). We note that the larger the log fold difference between comparisons the more the log fold change is underestimated. The variability of the fold change estimates of the spiked UPS proteins is very similar between all methods. Hence, the msqrob’s ridge penalisation only seems to affect the fold changes estimates of non-DA proteins and leaves those of DA proteins largely unaffected, which is a desirable property.

**Figure 4:**
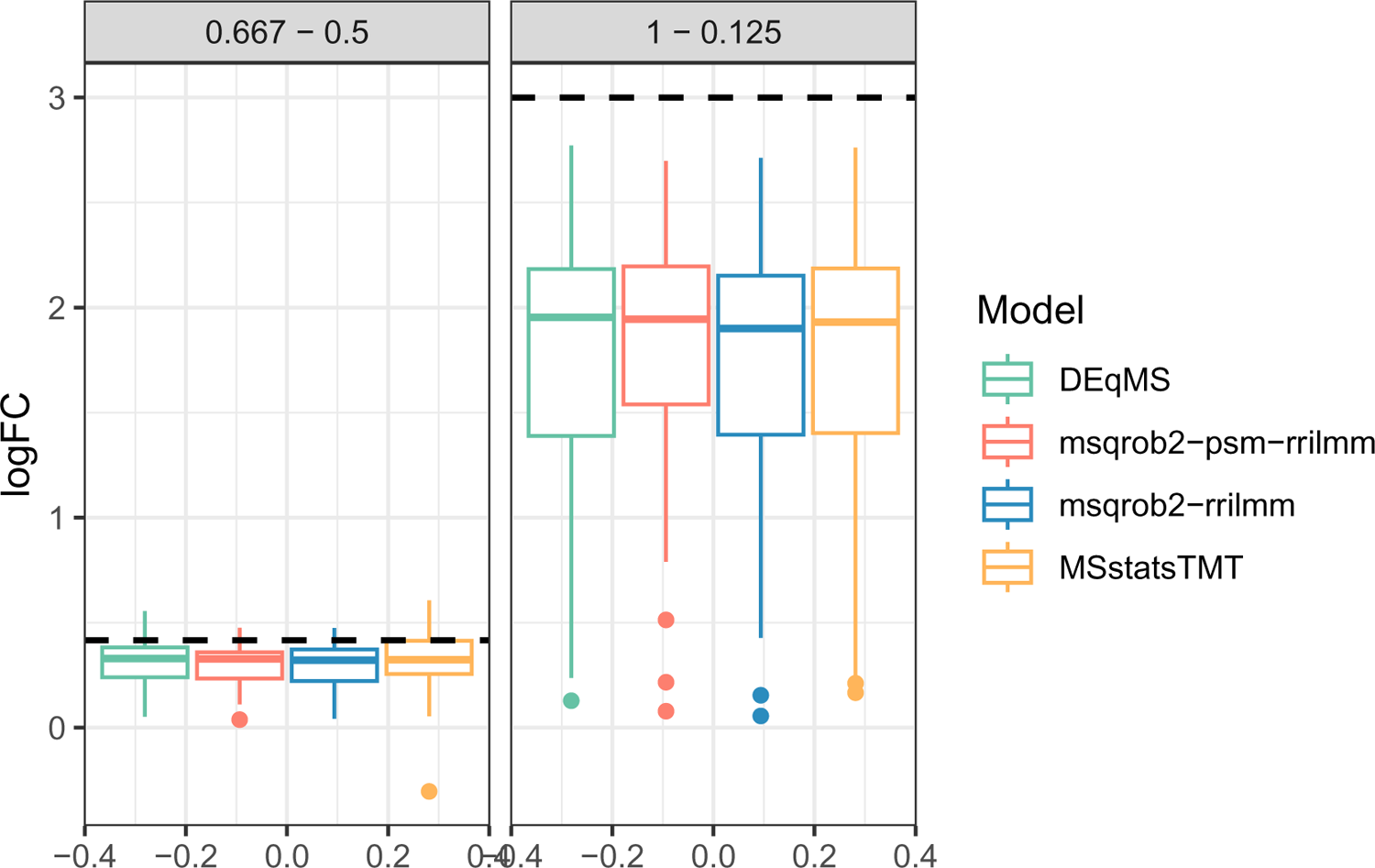
Boxplots showing the logFC distribution of the spiked-in proteins for the comparisons with the highest and lowest fold change between spike-in concentrations estimated by DEqMS, MSstatsTMT and the msqrob2TMT workflows. Black dotted line is the true logFC of the comparison.

Figure 5 shows that the MSstatsTMT and msqrob2TMT models produce fairly uniform p-value distributions for non spiked-in proteins. DEqMS shows a slight inflation at low p-values. The ridge regression p-values are slightly over-conservative and due to the shrinkage of the fold changes for the non spiked proteins there is a spike of p-values equal to 1.

**Figure 5:**
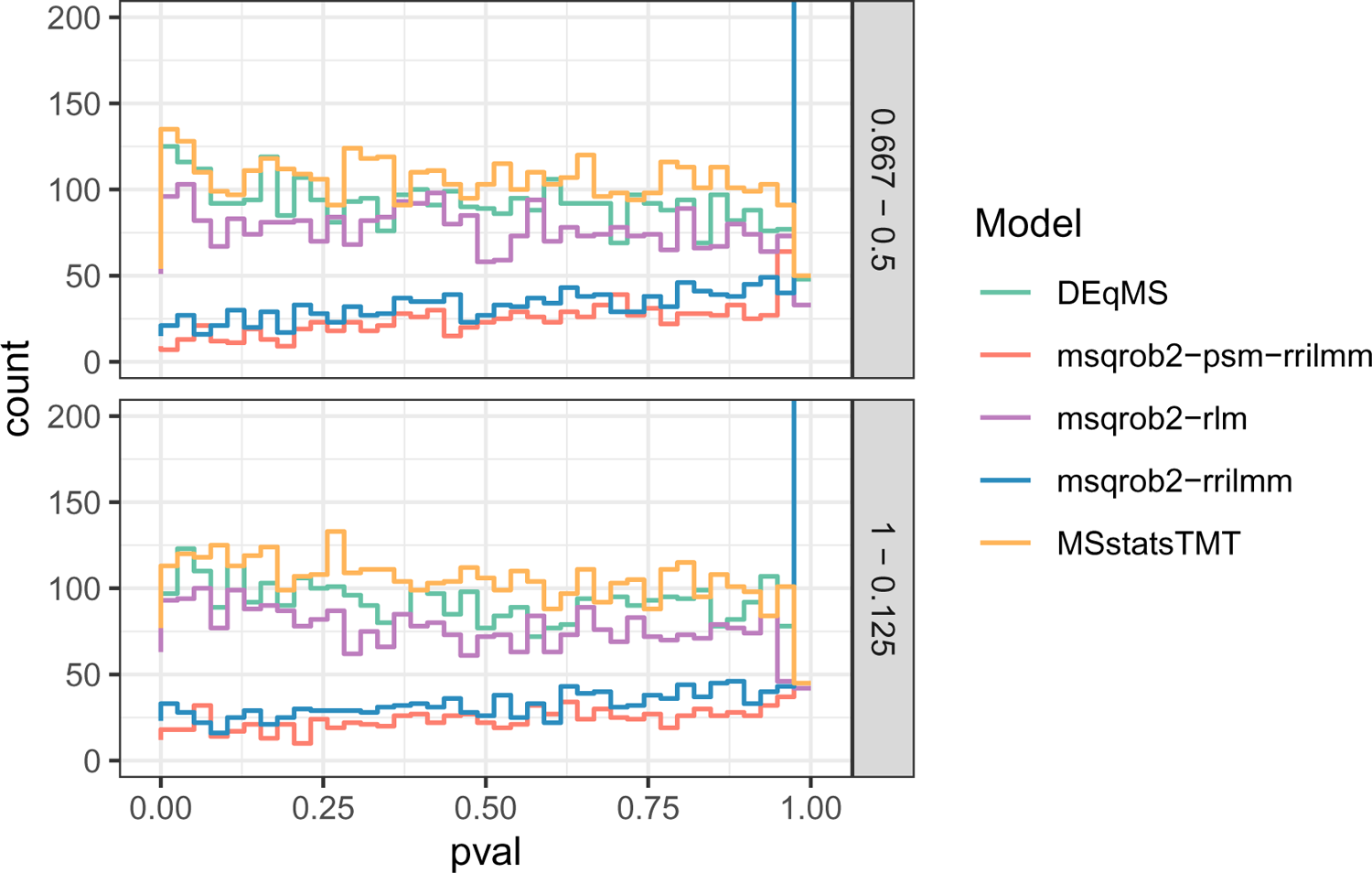
Histogram of the p-values from the non spiked-in proteins for the DEqMS, msqrob2TMT and MSstatsTMT workflows. Two comparisons between spike-in dilutions illustrating the differences are shown.

#### 3.1.2 Impact of M-estimation and ridge penalisation

As opposed to MSstatsTMT, the parameter estimation in msqrob2 can be regularised using robust M-estimation and/or ridge penalisation. In Figure 6 and Figure 7 we assess the impact of robust ridge regression on the performance in both the protein and PSM models. The largest impact can be seen in the comparison 0.667 versus 0.5 comparison involving the smallest fold change. We observe a small performance gain of robust M-estimation compared to a msqrob2TMT linear mixed model workflow. Ridge penalisation has a larger impact on the performance and the combination of robust M-estimation and ridge penalisation induces the largest performance gain. In the 1 vs 0.125 dilution comparison, only marginal differences between the curves can be seen. Overall the PSM models also have a higher precision.

**Figure 6:**
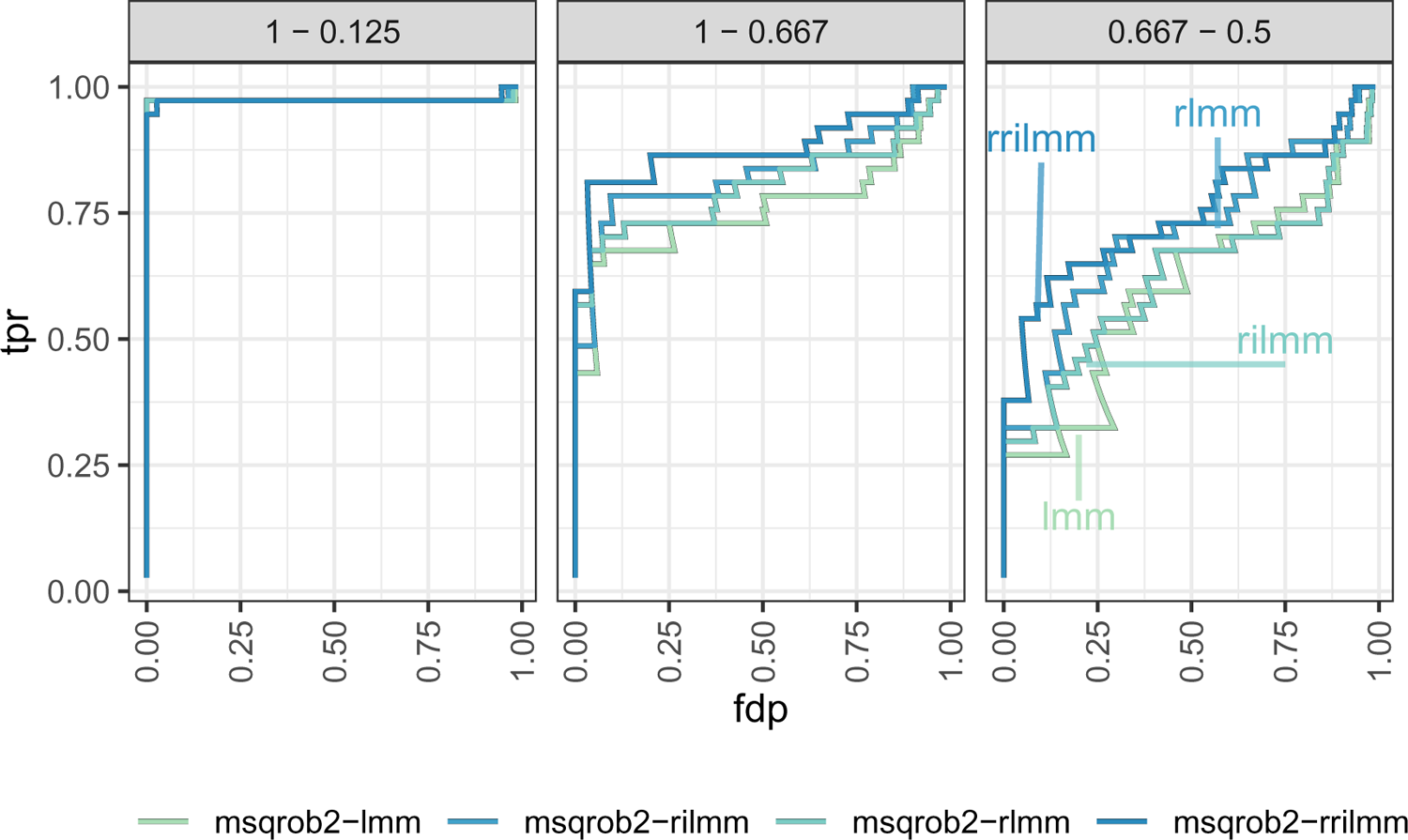
Impact of robust ridge regression on the performance. True positive rate (TRP) and false discovery proportion (FDP) is compared between msqrob2TMT protein level workflows without robust ridge regression, with only robust or ridge regression, and with both, for three effect sizes.

**Figure 7:**
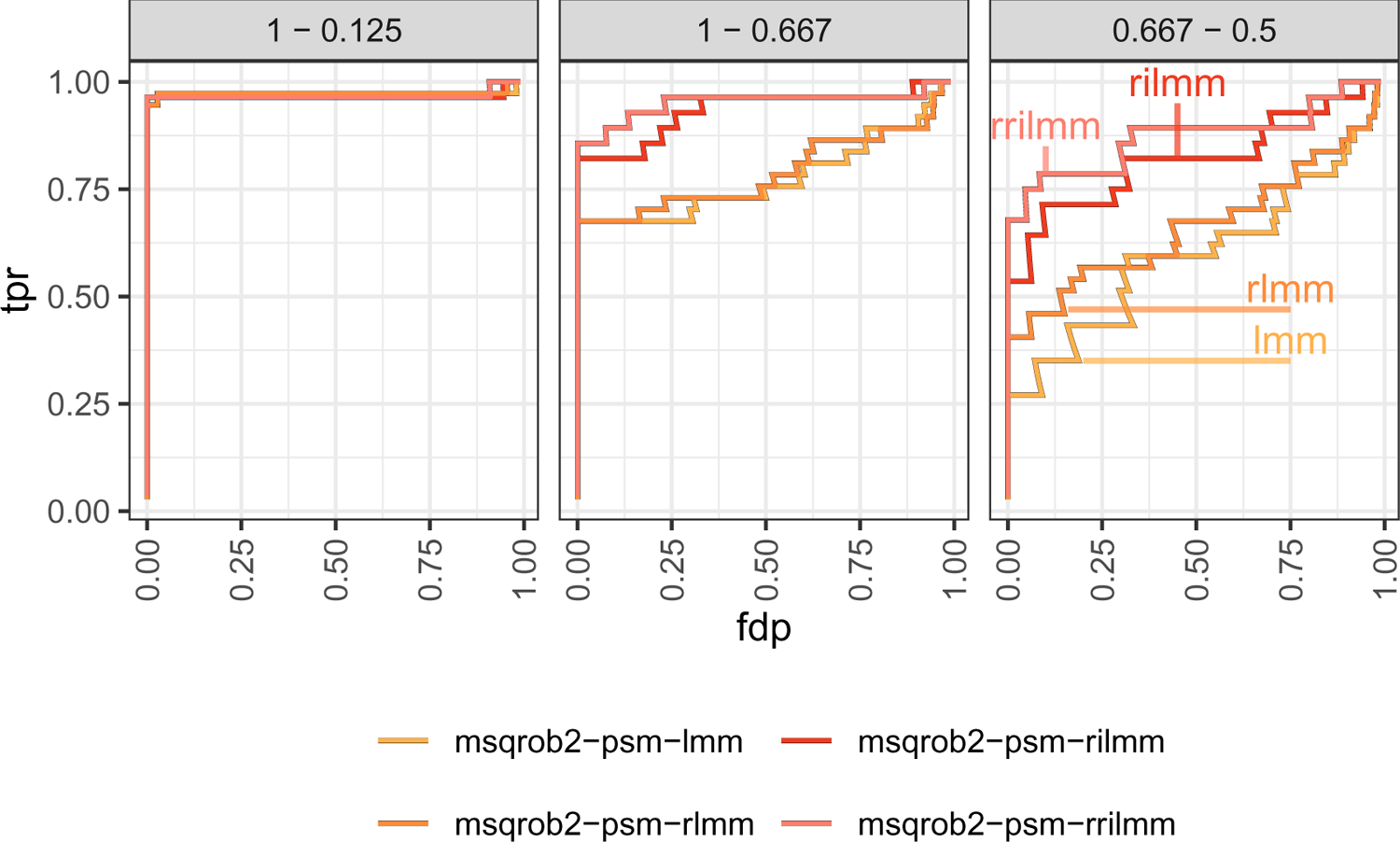
Impact of robust ridge regression on the performance. True positive rate (TRP) and false discovery proportion (FDP) is compared between msqrob2TMT PSM level workflows without robust ridge regression, with only robust or ridge regression, and with both, for three effect sizes.

Table 2 and Table 3 show the number of true and false positives at 0.05 FDR adjusted p-value cutoff for the protein and PSM models. Ridge regression decreases the number of true positives for comparison 1 vs 0.667 but it decreases the false positives in the largest comparison while keeping the number of true positives equal. Using robust regression increases the true positives while also increasing the number of false positives. The combination of both increases the true positives and reduces the number of false positives while controlling a false discovery proportion below 5%.

**Table 2:**
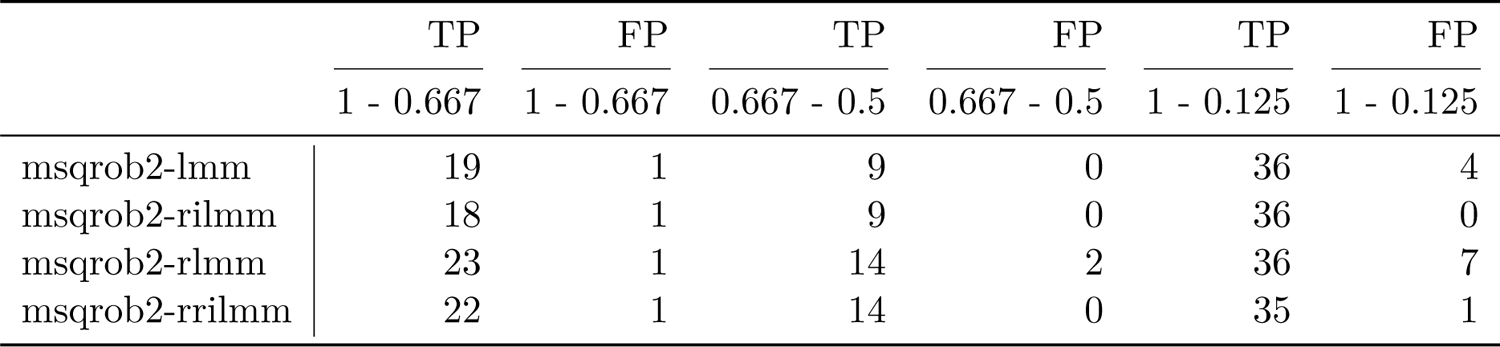
Impact of robust ridge regression on the performance. The number of true positives at 0.05 FDR adjusted p-value cutoff is compared between msqrob2TMT protein level workflows without robust ridge regression, with only robust or ridge regression and with both. Three spike-in concentration comparisons illustrating the differences are shown.

**Table 3:**
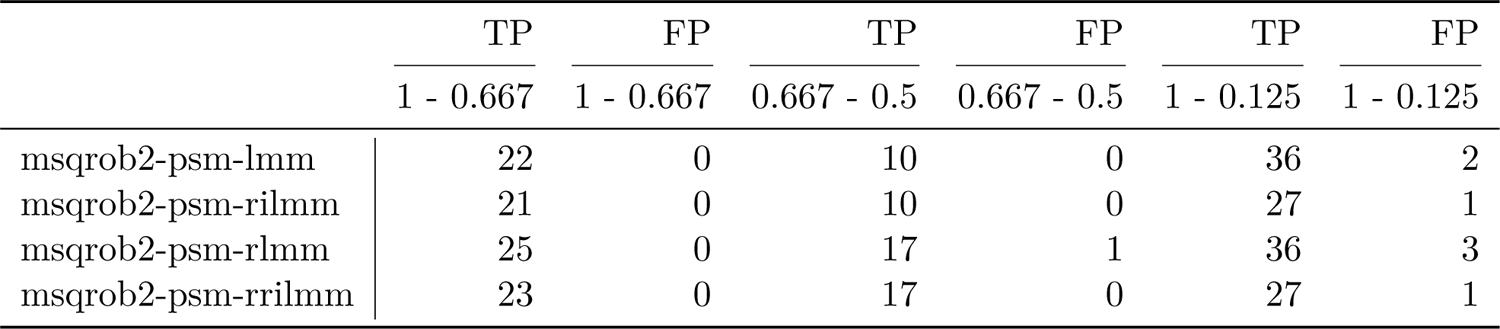
Impact of robust ridge regression on the performance. The number of true positives at 0.05 FDR adjusted p-value cutoff is compared between msqrob2TMT PSM level workflows without robust ridge regression, with only robust or ridge regression and with both. Three spike-in concentration comparisons illustrating the differences are shown.

#### 3.1.3 Impact of imputation and reference normalisation

Note, that MSstatsTMT, DEqMS and the msqrob2TMT based workflows also differ in the preprocessing, model estimation and hypothesis testing steps. We, therefore, have run DEqMS median sweep summarisation, MSstats pre-processing and summarization with and without imputation, and inferred on differential abundance with msqrob2-rrilmm.

Figure 8 panel A confirms that, the suggested msqrob2TMT workflow (linear mixed model using robust ridge regression) has superior performance. Performing the reference normalisation seems to negatively impact the performance, while imputation does not seem to have a definitive positive impact in this dataset. Note that, msqrob2TMT without robust ridge and with imputation and reference normalisation boils down to the MSstatsTMT workflow. Interestingly, however, Figure 8 panel B illustrates that the results of this msqrob2TMT workflow is still slightly better than those of MSstatsTMT. This is due to the choice in MSstatsTMT to reduce the model complexity automatically when the full linear mixed model cannot be fitted. Indeed, msqrob2TMT returned inference on 3626 proteins whereas MSstatsTMT could infer on 4212 proteins. Note, that fitting reduced models does not seem to improve the performance. With MSstatsTMT it is no longer clear for the user which proteins are fitted with random effects and for which proteins the random effects were dropped. For some proteins a reduced model is appropriate, but for others it might lead to ambiguous fits for which the hierarchical correlation structure is not properly addressed. With msqrob2 we have chosen to return a fit error if the protein cannot be estimated with the model that is specified by the user. We feel that the intervention of a skilled data analyst is needed to reduce the model for these proteins, which can be done by refitting the proteins with msqrob2.

**Figure 8:**
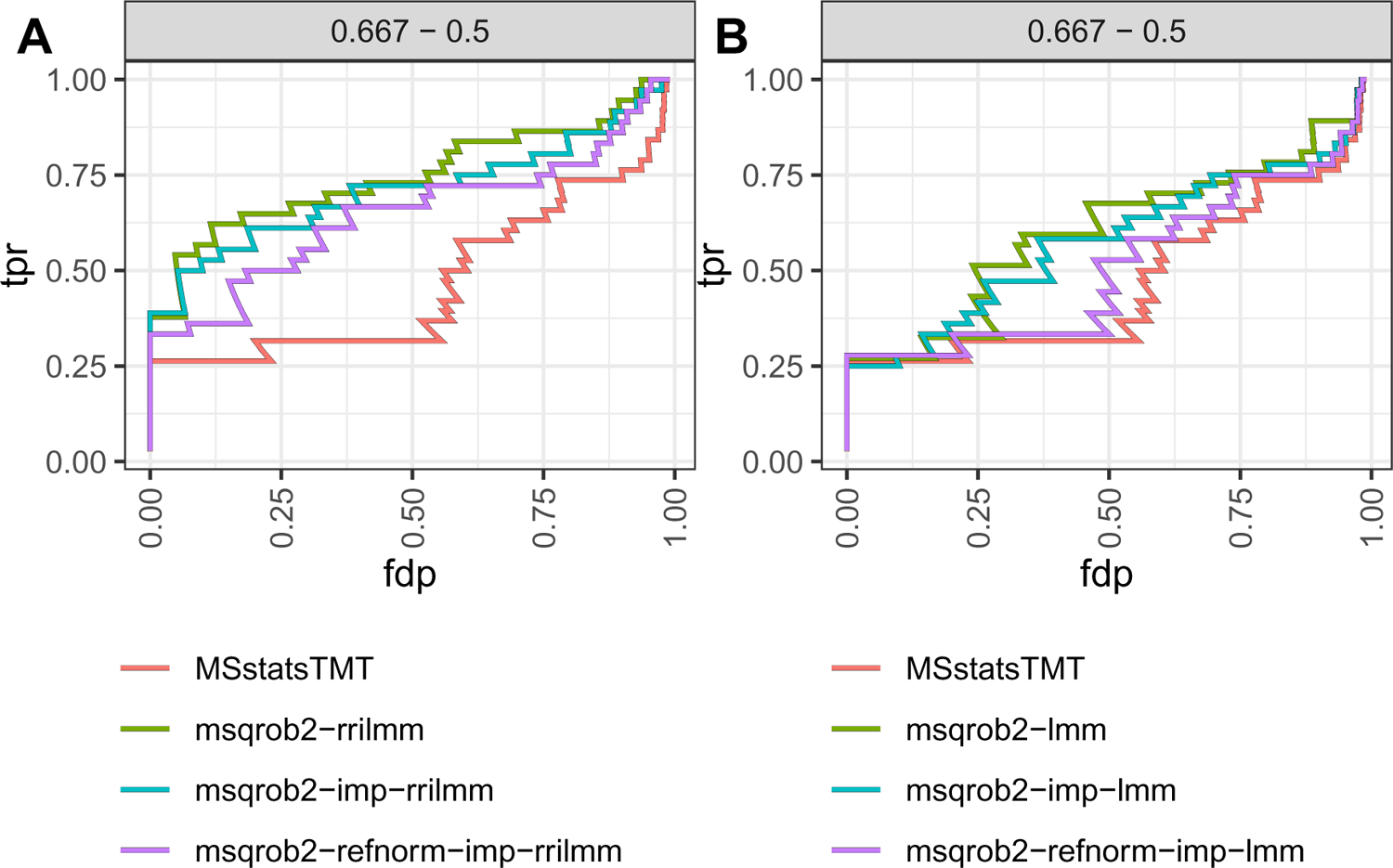
True positive rate (tpr) - false discovery proportion (fdp) plots for the msqrob2TMT workflows with imputation and reference normalisation and the MSstatsTMT workflow. Plot A shows the robust ridge linear mixed models and plot B shows the linear mixed models. refnorm refers to reference normalisation and imp refers to imputation, which is performed by using the accelerated time failure model. Results for all spike-in dilution comparisons can be found in Supplementary Figure: 2 and Supplementary Figure: 3.

### 3.2 Mouse study

The mouse dataset uses an unbalanced design where the mice have been fed a low fat (LF) or a high fat (HF) diet for a short or a long period. We will focus of the effect of the diet and duration on the proteome of adipocyte cells. Note, that MSstatsTMT cannot accommodate for designs with multiple factors and/or interactions. We therefore chose to encode the treatment effect using one factor with a level for each diet × duration combination.

Table 4 shows the number of differential proteins at 0.05 p-value cutoff for the unbalanced mouse dataset. The PSM model returns the most differential proteins. As we do not have the ground truth we are not able to verify if these proteins are truly differentially abundant.

**Table 4:** Table showing the number of differential proteins at 0.05 FDR adjusted p-value cutoff for the msqrob2TMT and MSstatsTMT workflows. Comparison 1 is the comparison between LongHF and LongLF, 2 is between LongHF and ShortHF, 3 is between LongLF and ShortLF, and 4 is between ShortHF and ShortLF.

|  | Comparison |  |  |  |
| --- | --- | --- | --- | --- |
|  | 1 | 2 | 3 | 4 |
| MSstatsTMT | 9 | 63 | 8 | 5 |
| msqrob2-psm-rrlmm | 37 | 130 | 86 | 96 |
| msqrob2-rrlmm | 8 | 49 | 23 | 38 |

Table 4 shows that the PSM level workflow msqrob2-psm-rrilmm consistently reports 2-4 times more DA proteins than MSstatsTMT and our protein level msqrob2-rrilmm workflow in all comparisons. MSstatsTMT reports 1 and 14 proteins more as DA than msqrob2-rrilmm in the first two comparisons. Conversely, in the third and fourth comparisons, MSstatsTMT reports 15 and 33 DA proteins less than msqrob2-rrilmm. The upset plots (Supplementary Figures: 5, 6, 7, 8) show that the majority of proteins that are reported by msqrob2-rrilmm are also picked up by the msqrob2-psm-rrilmm. All 8 and 5 DA proteins that were reported by MSstatsTMT in the third and fourth comparisons, were also picked up by protein level msqrob2-rrilmm workflow.

Taking closer look at the 6 unique DA proteins from comparison 1 (see Supplementary Figure 5) we find that two proteins Q3TBD2 and Q810U2 are not found in the msqrob2 models. These proteins were only found in one mixture, which implies that the full model could not be fitted. Our msqrob2TMT workflows therefore returned a fit error for these proteins, while MSstatsTMT automatically reduces the model to a linear model. Protein Q99JB8 has the biggest difference in adjusted p-values between the methods; 0.42, 0.21 and 0.01 for msqrob2-psm-rrilmm, msqrob2-rrilmm and MSstatsTMT, respectively. This is due to the large amount of missing data for this protein, which leads to large differences in the eventual logFC estimates; −0.46, −0.7 and −1.52, respectively. The other proteins Q99JW5, Q62470, and Q99N69 all have similar logFC’s and adjusted p-values for the different methods. However, they are not significant for the msqrob2TMT models but are close to the 0.05 adjusted p-value threshold.

The proteins that were reported DA by MSstats and not by our summarised msqrob2-rrilmm workflow in comparison 2 are due to similar reasons: 6 proteins were only picked up in one mixture, 4 due to missingness leading to differences in the log2 fold changes and 16 due to adjusted p-values that were just larger than the 5% threshold.

We also conducted a gene-set enrichment analysis for the comparison of longHF and shortHF diets as this comparison returned the largest lists of DA proteins (Table 5).

**Table 5:** Gene ontology tables showing the significant KEGG pathways. Background used all mmusculus proteins. Results shown for comparision between the longHF and shortHF diets.

|  | term_id | p_value |
| --- | --- | --- |
| msqrob rilmm psm |  |  |
| Complement and coagulation cascades | KEGG:04610 | 4.482777e-13 |
| Biosynthesis of amino acids | KEGG:01230 | 2.222546e-03 |
| Staphylococcus aureus infection | KEGG:05150 | 2.949620e-03 |
| Metabolic pathways | KEGG:01100 | 6.478388e-03 |
| Pentose phosphate pathway | KEGG:00030 | 1.021939e-02 |
| Glycolysis / Gluconeogenesis | KEGG:00010 | 1.173015e-02 |
| Alanine, aspartate and glutamate metabolism | KEGG:00250 | 1.589693e-02 |
| Carbon metabolism | KEGG:01200 | 2.459682e-02 |
| msqrob rilmm |  |  |
| Complement and coagulation cascades | KEGG:04610 | 1.704394e-05 |
| Metabolic pathways | KEGG:01100 | 4.751297e-03 |
| Pentose phosphate pathway | KEGG:00030 | 8.815811e-03 |
| Carbon metabolism | KEGG:01200 | 2.718292e-02 |
| MSstatsTMT |  |  |
| Carbon metabolism | KEGG:01200 | 3.966898e-03 |
| Glycolysis / Gluconeogenesis | KEGG:00010 | 5.516530e-03 |
| Biosynthesis of amino acids | KEGG:01230 | 1.059700e-02 |
| Metabolic pathways | KEGG:01100 | 1.108744e-02 |
| Pentose phosphate pathway | KEGG:00030 | 1.541934e-02 |
| PPAR signaling pathway | KEGG:03320 | 1.764945e-02 |
| Complement and coagulation cascades | KEGG:04610 | 2.089660e-02 |
| Glycine, serine and threonine metabolism | KEGG:00260 | 2.326707e-02 |

The gene set enrichment analysis can be performed against a background consisting of the entire mouse proteome or against the proteins that were picked up in the study. The latter is more convenient. Indeed, the proteins are isolated from mouse adipose tissue so we can expect an overrepresentation of adipose tissue related proteins. With this analysis only the “complement and coagulation cascade KEGG pathway” (p-value 7.4*10^-6^) was significant with the msqrob2TMT psm workflow while none of the KEGG pathways were returned by the MSstats and the msqrob2TMT protein level workflows. Therefore we also ran the gene set enrichment analysis against the entire mouse background proteome for a more broad comparison.

For the gene-set enrichment analysis against the entire mouse proteome, the results between the workflows are fairly similar. The largest difference is found in proteins related to complement and coagulation cascades. More proteins related to the coagulation pathway are found in the PSM workflow, these results are also verified in literature. These workflows are based on a 5% FDR cutoff, while the original paper uses a 10% FDR cutoff.

Validity of the p-values was verified in a mock analysis. Here we added an additional mock variable that splits the samples at random in two groups per run. Hence, we know that all proteins are not DA between both mock groups, so their p-values should be uniform. The mock analysis could only be done in msqrob2TMT workflows as it is not possible to analyze a two factor design with main effects with MSstatsTMT. Indeed, the treatment effect for this design can no longer be encoded using a single factor with multiple levels.

Figure 9 shows that the p-value distributions for the robust ridge are uniform with a spike on 1. This pattern was also observed in the spike-in study and is due to the shrinkage of the model parameters towards zero. The mock analysis indicates that the additional proteins that are reported to be significant with the psm-level model do not seem to be a consequence of asymptotic inference that is too liberal.

**Figure 9:**
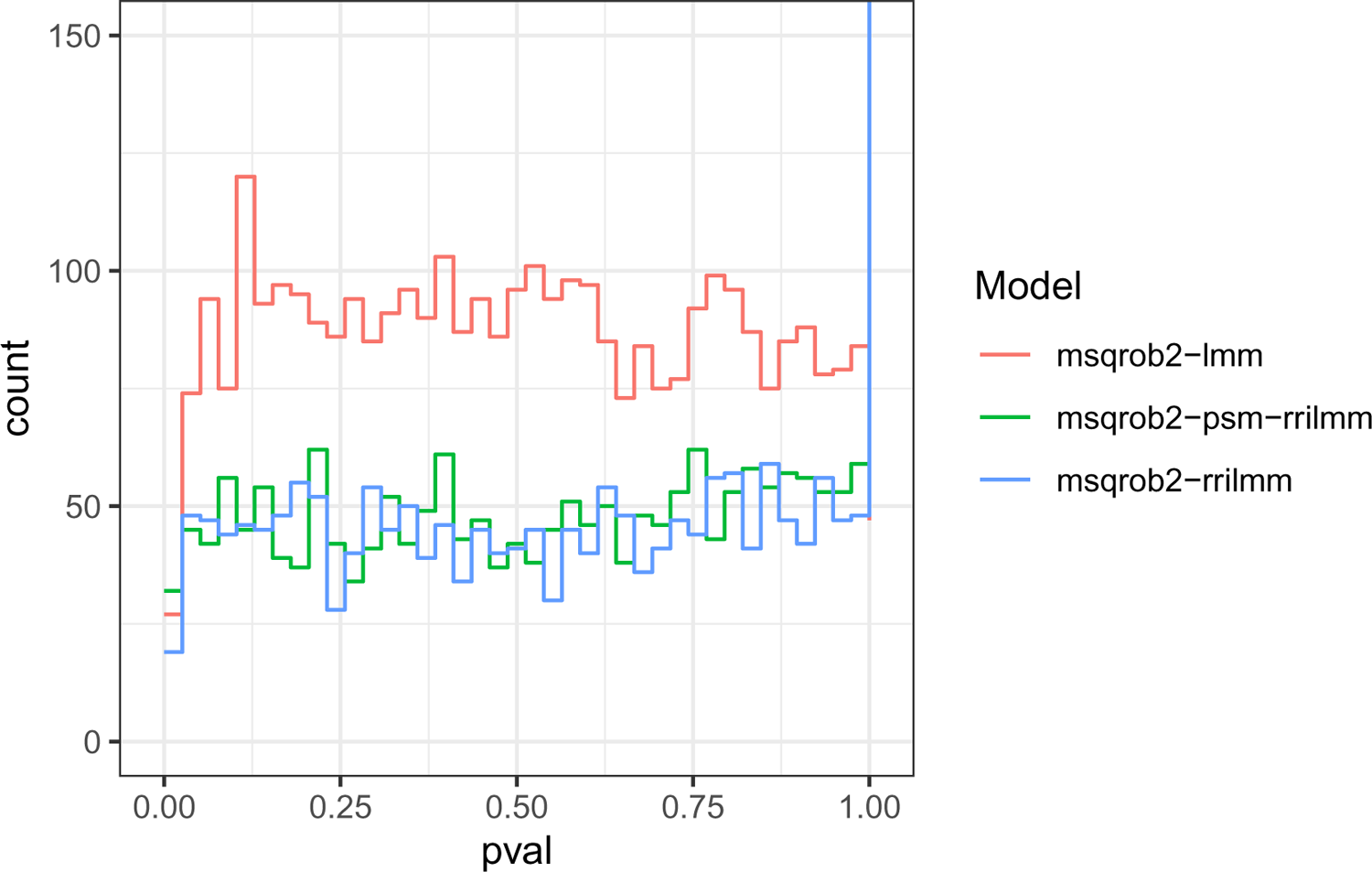
Histogram of the p-values from the mock analysis of the unbalanced mouse dataset. Msqrob2TMT workflows for both aggregated protein intensities and the PSM models are shown with and without robust ridge regression.”

## 4 Discussion

In this contribution we introduced novel workflows in the msqrob2 universe, designed for performing differential abundance analysis for labelled mass spectrometry based proteomics experiments. These workflows capitalise on msqrob2’s flexible robust ridge regression framework to improve the performance as compared to the state-of-the-art tools MSstatsTMT that uses a standard linear mixed model. In our newest 1.10 release we have unlocked full compatibility with the lme4 R package for mixed models, unlocking more complex designs that can use both feature level and sample level covariates. This means that our workflows can now address the complex hierarchical correlation structure of the data from labelled MS experiments, using both feature level and sample level covariates, and allow the user to analyze data from experiments with complex layouts that fit within the general class of linear mixed models, whereas MSstatsTMT only allows the inclusion of one additional factor on top of a random run and mixture effect. In the analysis of the spike-in benchmark dataset we clearly demonstrated that our novel msqrob2TMT workflows improve upon the state-of-the art methods in terms of sensitivity, specificity and FDR control. We showed that parameter estimation procedure is crucial for the performance in a DA workflow. msqrob2 extends the linear mixed model to robustify it against outliers and to improve uncertainty estimation, which lead to considerable performance gains and this especially in the more challenging comparisons involving low fold changes between the spike-in proteins (Figure 2). Another crucial component for the performance in a labelled MS-based DA workflow is the preprocessing. The first type of such preprocessing is normalisation, which can have a large impact on DE analysis. The second type of preprocessing is imputation of missing values. Different sources of missingness occur, which can lead to suboptimal results when the data are imputed under the assumption of missingness by low abundance. The robust modelling in msqrob2, however, can safely omit imputation altogether (Figure 8). Our workflows are also modular and provide the user with the flexibility to use custom pre-processing steps, which renders it future proof when novel and more performant normalisation, summarization and imputation procedures become available. Besides the fitting procedure, MSstatsTMT and msqrob2 also differ in their calculation of the degrees of freedom. Indeed, there is no consensus on which approach to apply for the calculation of the degrees of freedom for mixed models. msqrob2 use the residual effective degrees of freedom approach while MSstatsTMT uses the Satterthwaite approximation. Both tools use an empirical Bayes variance estimator using the “squeezevar” function from limma. However, MSstatsTMT adds the prior degrees of freedom from the empirical Bayes variance estimator for the residual variance to the degrees of freedom of the variance estimator for the contrast obtained with the Satterthwaite approximation, and it is unclear if that is theoretically justi-fied. Indeed, the prior degrees of freedom only involve one source of variability: the residual variance while the variance estimator of the contrast involves both the residual variance and random effect variances, and for the latter no empirical Bayes shrinkage across proteins has been applied.

A large difference between the msqrob2 and MSstatsTMT packages is that MSstatsTMT tries to fit as many proteins as possible and automate the model fitting. This has the advantage of being easier to use, however it does lead to fitting proteins from the same dataset with different models, i.e. by combining linear models with mixed models, which may not be appropriate for the analysis. Although it is possible to retrieve the models, and check their validity, a standard user is not fully aware of these issues, and the subtleties of interpretation that these require. In contrast, our msqrob2TMT workflows emphasise transparency and reproducibility. So we chose not to reduce the models automatically when the model that is specified by the user cannot be fitted. Instead, we report a fit-error and the data analyst can then refit the specific proteins with an appropriate model.

Our analyses also illustrate the need of good benchmark datasets in the proteomics field. Indeed, the spike-in dataset has its limitations as the results heavily depend on the pre-processing of the UPS proteins. The background proteins originate from HeLa cells, which also contain UPS proteins. The background UPS proteins and the spiked-in UPS proteins differ in metabolic labelling, so we should be able to distinguish them. We used searched PSM-level data from mascot that was provided by the MSstatsTMT authors. In their workflow, two mascot identification nodes were used. In one node they searched the SwissProt database for proteins with static modifications related to the metabolic labelling, in the other node they searched the Sigma_UPS protein database without these static modifications. Ideally this should seperate the spiked-in UPS proteins and the UPS proteins from the HeLa cells, however this was not the case. When closely examining the results, we found spectra that match perfectly with a DA pattern expected for spike-in proteins, whilst not being tagged as UPS and vice versa. We therefore performed additional pre-processing steps in order to remove these ambiguous proteins. It is, however, possible that some spectra are still mislabelled. This can lead to two issues: on the one hand excess false positives originating from spike-in proteins that were erroneously not tagged as spiked-in UPS proteins. On the other hand, a lower sensitivity as including spectra from wrongly labelled background UPS proteins reduces the log fold change estimates and increases the standard errors of the estimates, leading to spike-in UPS proteins not being picked up as significant. When performing the analysis without the extra pre-processing, we found more false positives, however, these were background proteins that were not tagged as spike-in UPS proteins, but that clearly showed a DA pattern as expected for spike-in UPS proteins. Another issue related to the setup of the benchmark study is that all spike-in proteins are DA in the same direction. This can lead to issues with the channel normalisation, as other proteins can be biased downwards due to ion suppression effects in high spike-in samples. This is evidenced in the results. Indeed, more false positives are found in comparisons involving samples with a larger difference in spike-in dilution (see Supplementary Table: 2). Hence, to advance the field benchmark datasets are required with more complex DA patterns where some proteins are upregulated, and others are downregulated in each spike-in condition.

Overall, we have shown that our msqrob2TMT workflow is a sensitive and robust approach compared to the state-of-the-art, while providing good FDR control. Our modular implementation offers our users full flexibility with respect to the search engine and pre-processing steps, while still offering a comprehensive, transparent, and reproducible workflow that covers the entire differential proteomics analysis.

## 7 Supplementary Materials

### 7.1 Spike-In Dataset

**Supplementary Figure 1:**
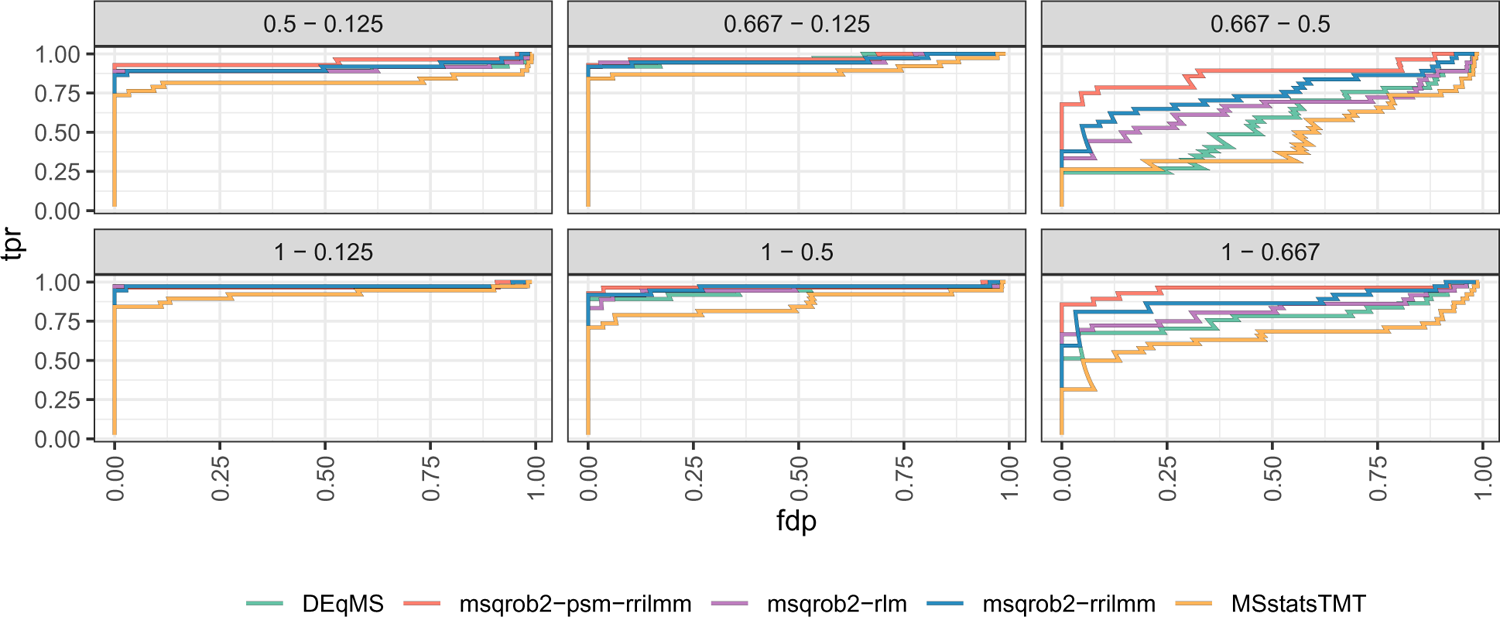
Figure showing the true positive rate in relation to the false discovery proportion for the DEqMS, msqrob2TMT and MSstatsTMT workflows

**Supplementary Table 1:** Table showing the number of true positives (TP) and false positives (FP) at 0.05 FDR adjusted p-value cutoff for the DEqMS, msqrob2 and MSstatsTMT workflows.

(a)
|  | TP | FP | TP | FP | TP | FP |
| --- | --- | --- | --- | --- | --- | --- |
|  | 1 - 0.667 | 1 - 0.667 | 0.667 - 0.5 | 0.667 - 0.5 | 1 - 0.125 | 1 - 0.125 |
| DEqMS | 21 | 1 | 9 | 0 | 36 | 5 |
| MSstatsTMT | 16 | 1 | 5 | 0 | 34 | 5 |
| msqrob2-psm-rrilmm | 23 | 0 | 17 | 0 | 27 | 1 |
| msqrob2-rlm | 21 | 0 | 12 | 1 | 35 | 5 |
| msqrob2-rrilmm | 22 | 1 | 14 | 0 | 35 | 1 |

(b)
|  | TP | FP | TP | FP | TP | FP |
| --- | --- | --- | --- | --- | --- | --- |
|  | 0.5 - 0.125 | 0.5 - 0.125 | 0.667 - 0.125 | 0.667 - 0.125 | 1 - 0.5 | 1 - 0.5 |
| DEqMS | 33 | 2 | 34 | 3 | 33 | 5 |
| MSstatsTMT | 31 | 4 | 32 | 2 | 28 | 1 |
| msqrob2-psm-rrilmm | 26 | 0 | 26 | 1 | 26 | 0 |
| msqrob2-rlm | 32 | 3 | 33 | 1 | 32 | 2 |
| msqrob2-rrilmm | 32 | 0 | 34 | 2 | 32 | 0 |

#### 7.1.1 msqrob2TMT workflow with MSstatsTMT imputation

**Supplementary Figure 2:**
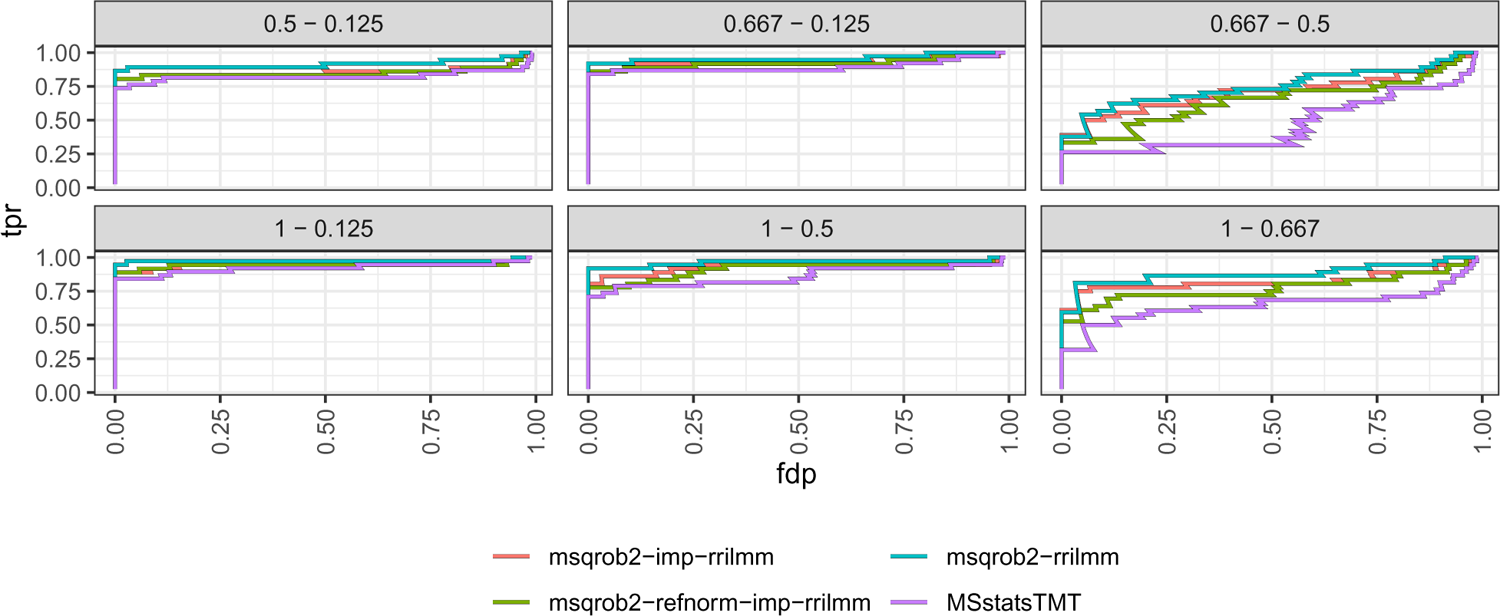
Figure showing the true positive rate in relation to the false discovery proportion for the msqrob2TMT workflows with imputation and reference normalisation and the MSstatsTMT workflow. refnorm refers to reference normalisation and imputation refers to the accelerated time failure model.

**Supplementary Figure 3:**
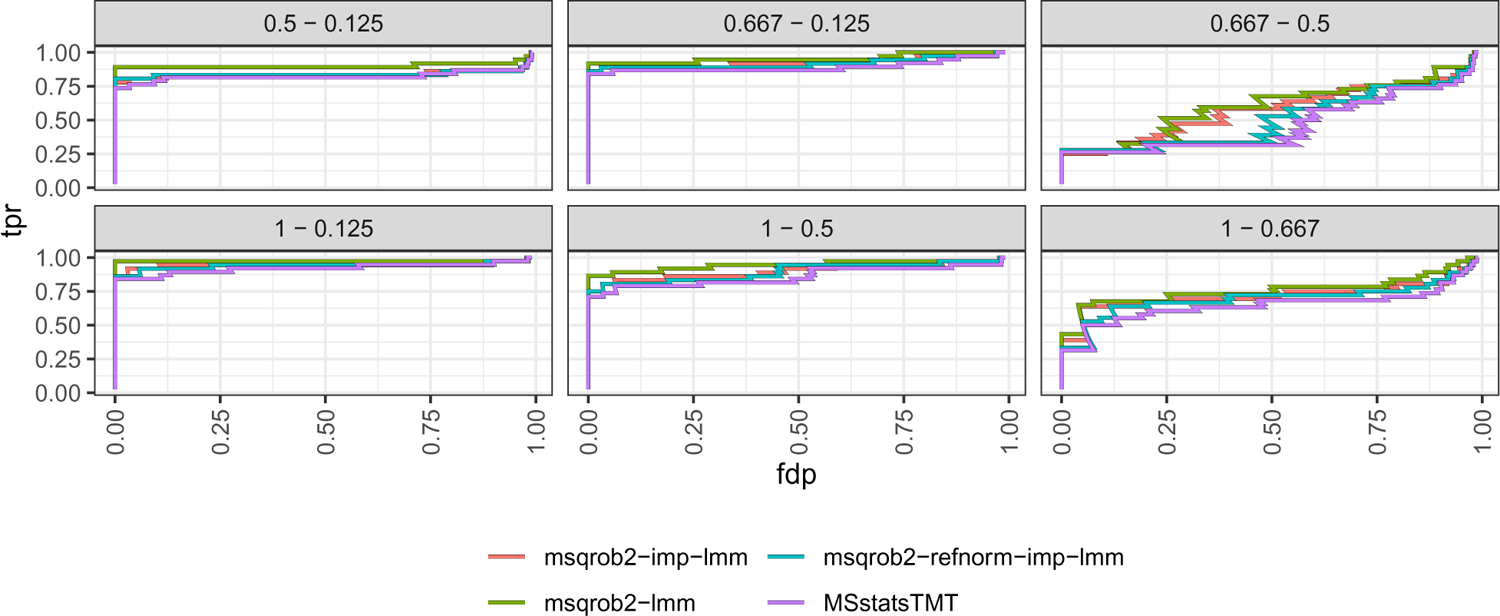
Figure showing the true positive rate in relation to the false discovery proportion for the msqrob2TMT workflows with imputation and reference normalisation and the MSstatsTMT workflow. refnorm refers to reference normalisation and imputation refers to the accelerated time failure model

#### 7.1.2 Dataset with technical replicates

**Supplementary Figure 4:**
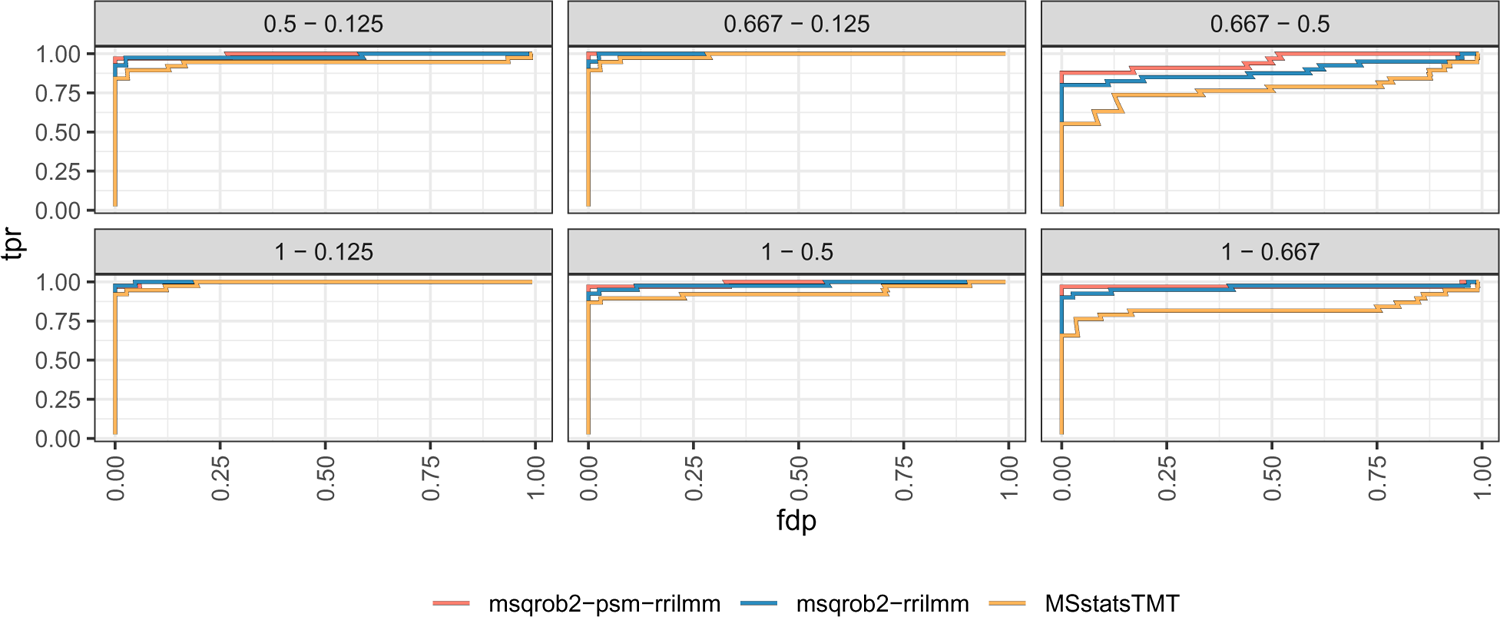
Figure showing the true positive rate in relation to the false discovery proportion for the msqrob2TMT and MSstatsTMT workflows using all the technical replicates in the dataset.

**Supplementary Table 2:** Table showing the number of true positives (TP) and false positives (FP) at 0.05 FDR adjusted p-value cutoff for the DEqMS, msqrob2TMT and MSstatsTMT workflows using all the technical replicates in the dataset.

|  | TP | FP | TP | FP | TP | FP |
| --- | --- | --- | --- | --- | --- | --- |
|  | 1 - 0.667 | 1 - 0.667 | 0.667 - 0.5 | 0.667 - 0.5 | 1 - 0.125 | 1 - 0.125 |
| MSstatsTMT | 29 | 1 | 19 | 0 | 36 | 5 |
| msqrob2-psm-rrilmm | 31 | 0 | 28 | 0 | 33 | 6 |
| msqrob2-rrilmm | 35 | 0 | 32 | 0 | 40 | 3 |

(b)
|  | TP | FP | TP | FP | TP | FP |
| --- | --- | --- | --- | --- | --- | --- |
|  | 0.5 - 0.125 | 0.5 - 0.125 | 0.667 - 0.125 | 0.667 - 0.125 | 1 - 0.5 | 1 - 0.5 |
| MSstatsTMT | 34 | 3 | 36 | 3 | 34 | 2 |
| msqrob2-psm-rrilmm | 32 | 0 | 33 | 2 | 32 | 0 |
| msqrob2-rrilmm | 39 | 1 | 40 | 1 | 37 | 1 |

### 7.2 Mouse Dataset

**Supplementary Figure 5:**
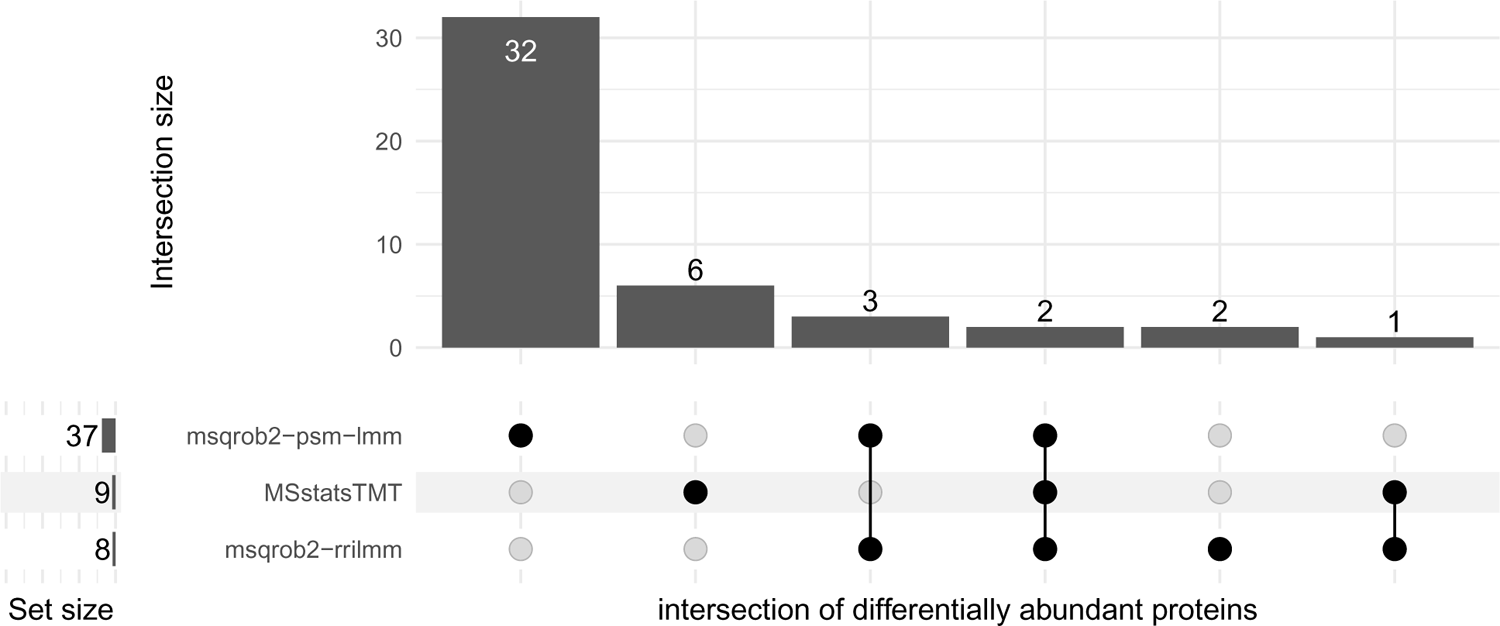
Upset plot showing the overlap in significant proteins between the MSstatsTMT and the msqrob2TMT workflows for the LongHF-LongLF comparison.

**Supplementary Figure 6:**
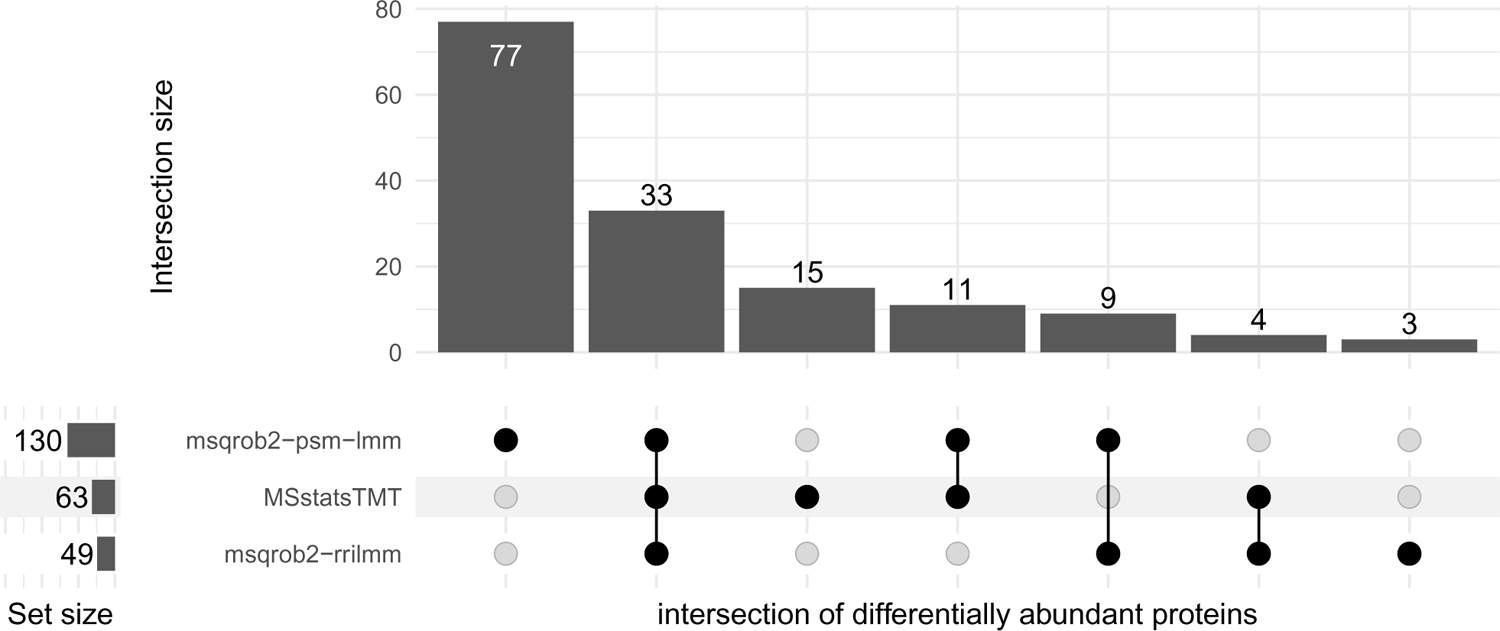
Upset plot showing the overlap in significant proteins between the MSstatsTMT and the msqrob2TMT workflows for the LongHF-LongLF comparison.

**Supplementary Figure 7:**
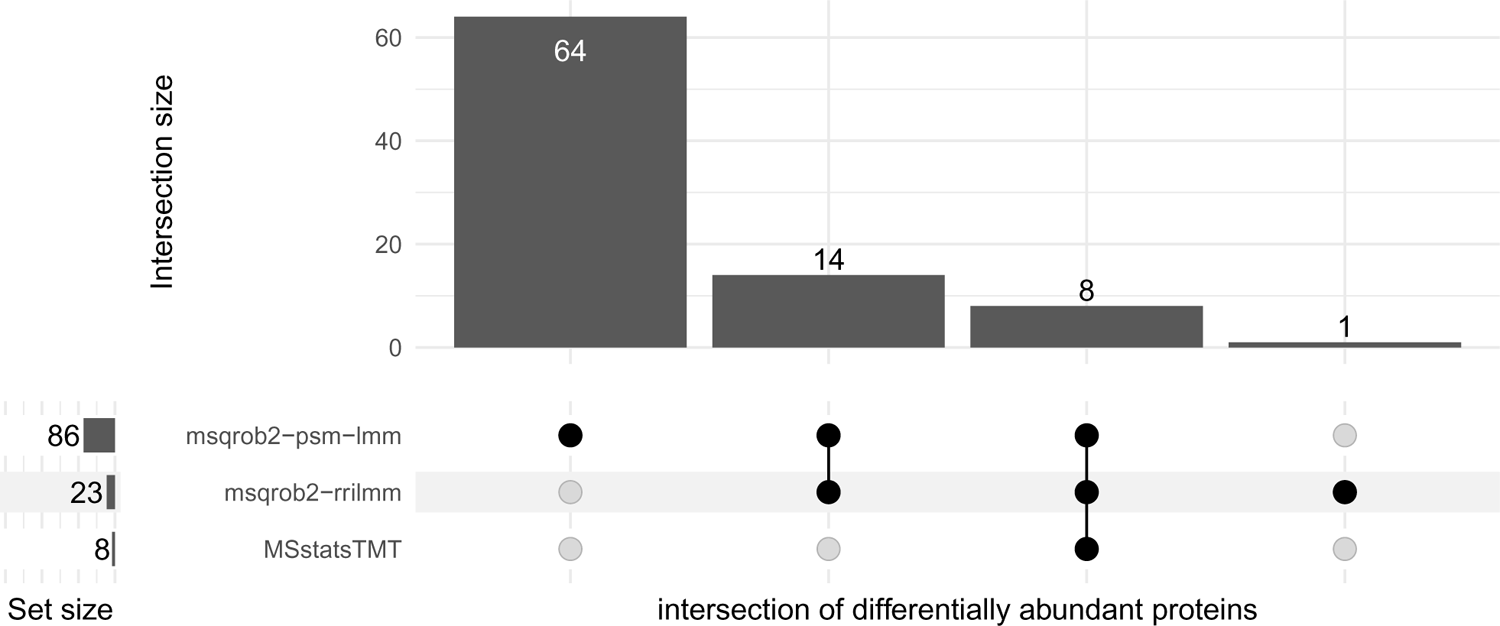
Upset plot showing the overlap in significant proteins between the MSstatsTMT and the msqrob2TMT workflows for the LongLF-ShortLF comparison.

**Supplementary Figure 8:**
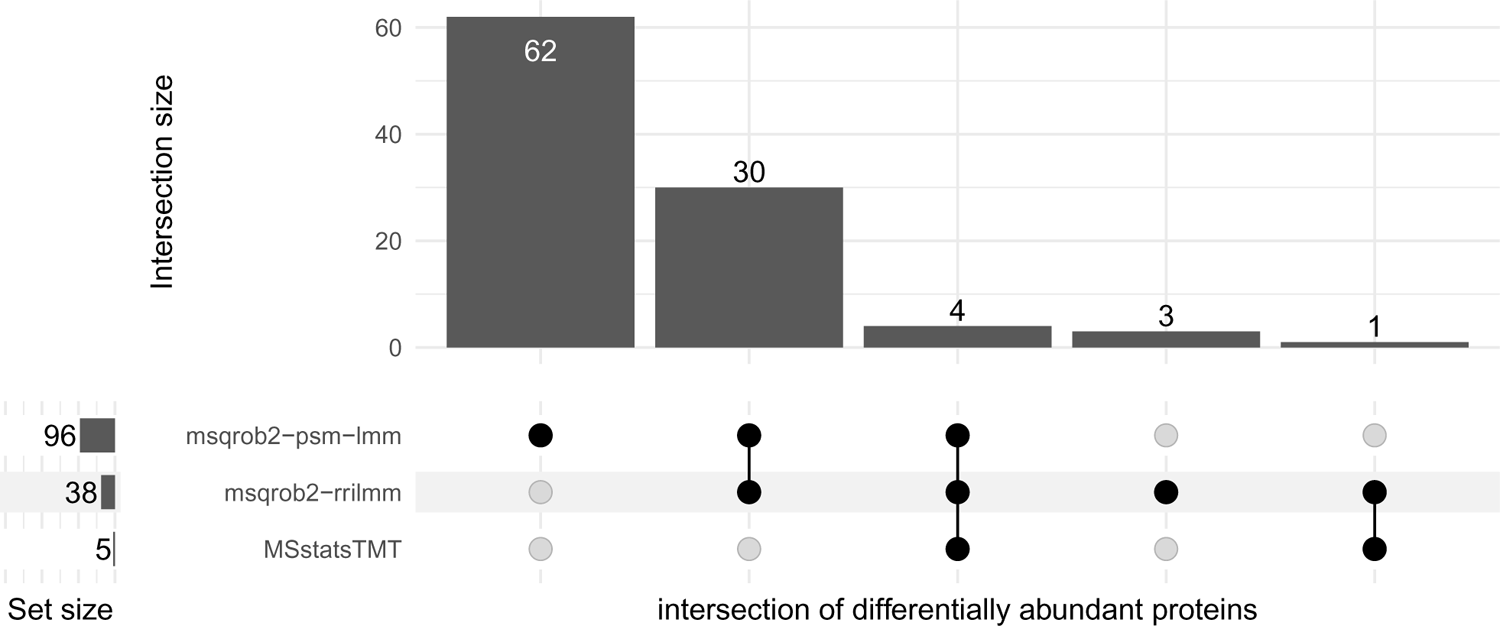
Upset plot showing the overlap in significant proteins between the MSstatsTMT and the msqrob2TMT workflows for the ShortHF-ShortLF comparison.

